# Systematic Engineering of Intra-Articular Drug Release Profiles Reveals a Key Determinant of Disease-Modifying Efficacy in Post-Traumatic Osteoarthritis

**DOI:** 10.64898/2026.05.30.728894

**Authors:** Jingjing Gao, Nutan Bhingaradiya, Ziting Judy Xia, Ryan Yip, Eli Weldon, Bou Chosson Leite Chilan, Nishkal Dhiraj Pisal, Swetharajan Gunasekar, Pranav Chandrasekar, Paula Oliva Ribas, Mahima Dewani, Christopher Jiang, Gopinathan Janarthanan, Alexandra Dolliver, Rachel Wai Chun Chu, Gunjan Malik, Sohyung Lee, Ranit Dutta, Sanjairaj Vijayavenkataraman, Jeffrey M. Karp, Joerg Ermann, Nitin Joshi

## Abstract

Post-traumatic osteoarthritis (PTOA) is a progressive joint disease for which no disease-modifying osteoarthritis drugs (DMOADs) have been approved. Although injectable drug delivery systems can prolong therapeutic retention within the joint, it remains unclear whether local drug release kinetics influence disease-modifying efficacy. Here, we developed a modular platform of injectable supramolecular hydrogels using biocompatible, generally recognized as safe (GRAS) amphiphilic molecules and systematically engineered a range of degradation and drug release profiles. Using the cathepsin-K inhibitor L-006235 as a model DMOAD, we generated hydrogels with distinct release kinetics and evaluated their therapeutic performance in PTOA. Hydrogels exhibiting slower degradation and more sustained drug release like Sucrose Stearate (SS hydrogel) showed prolonged intra-articular retention and improved therapeutic outcomes. In a destabilization of the medial meniscus (DMM) mouse model, sustained-release formulations significantly reduced cartilage degeneration, preserved aggrecan expression, improved joint histopathology, and enabled effective monthly dosing. In contrast, formulations with faster degradation and release kinetics required more frequent administration to achieve comparable benefits. To our knowledge, this is the first study to establish local drug release kinetics as a critical determinant of disease-modifying efficacy in PTOA. This work provides one of the clearest demonstrations to date that engineering intra-articular release kinetics, rather than merely prolonging residence time, can improve disease-modifying outcomes. Our findings establish local release kinetics as a key design parameter for osteoarthritis therapeutics and highlight the potential of tunable supramolecular hydrogels for long-acting drug delivery.

## Introduction

Post-traumatic osteoarthritis (PTOA) is a debilitating degenerative joint disease that develops following joint injuries, such as meniscus or ligament tears or damage to cartilage, which lead to progressive cartilage degradation, chronic pain, and functional impairment.^1^ Most available pharmacological treatments, such as nonsteroidal anti-inflammatory drugs (NSAIDs) and corticosteroids, only provide symptomatic relief without addressing the underlying disease mechanisms. To date, there are currently no FDA-approved disease-modifying osteoarthritis drugs (DMOADs) capable of halting or reversing PTOA progression.^2–5^

A central barrier to developing effective disease-modifying osteoarthritis drugs (DMOADs) lies in the rapid clearance of therapeutic agents from the joint space.^5^ This results in brief and subtherapeutic drug exposure at target tissues. Small-molecule inhibitors, for example, often exhibit intra-articular half-lives of less than two hours.^6^ To counter this limitation, a variety of local delivery systems, including nanoparticles, microparticles, and hydrogels have been developed to prolong residence time and achieve sustained release within the joint.^7–11^ While these platforms successfully extend drug exposure, the kinetics of local drug release, rather than duration alone, can critically shape pharmacodynamic responses and therapeutic outcomes. This principle has been demonstrated in other diseases, where precise tuning of local release rates has determined therapeutic efficacy. Yet, in osteoarthritis, the relationship between local release kinetics and disease modification remains largely unexplored. Understanding and engineering this relationship is essential to move beyond sustained exposure toward optimized temporal control of intra-articular drug delivery, an advance that could fundamentally redefine how DMOADs are designed and deployed to prevent or reverse joint degeneration.

We previously developed a mechanically resilient, self-assembling hydrogel composed of a low-molecular-weight gelator (LMWG) - triglycerol monostearate (TG-18) - that exhibits thixotropic behavior, allowing it to rapidly recover from mechanical loading conditions relevant to physically active human joints, thus exhibiting sustained release of cathepsin-K inhibitor L-006235, a proof-of-concept small-molecule DMOAD.^12^ A small volume (4 µL) of this injectable system enabled sustained intra-articular release for over two weeks in physically active mice and effectively prevented structural and functional progression of post-traumatic osteoarthritis. This hydrogel demonstrated superior performance compared to a previously reported hydrogel designed for intra-articular drug delivery, which, in our study, neither recovered its structure nor maintained drug release under mechanical loading.^13,14^ Here, we demonstrate that tuning the local release kinetics of L-006235 can influence its efficacy in PTOA treatment and further prolong its residence time (Figure 1A).

**Figure 1.**
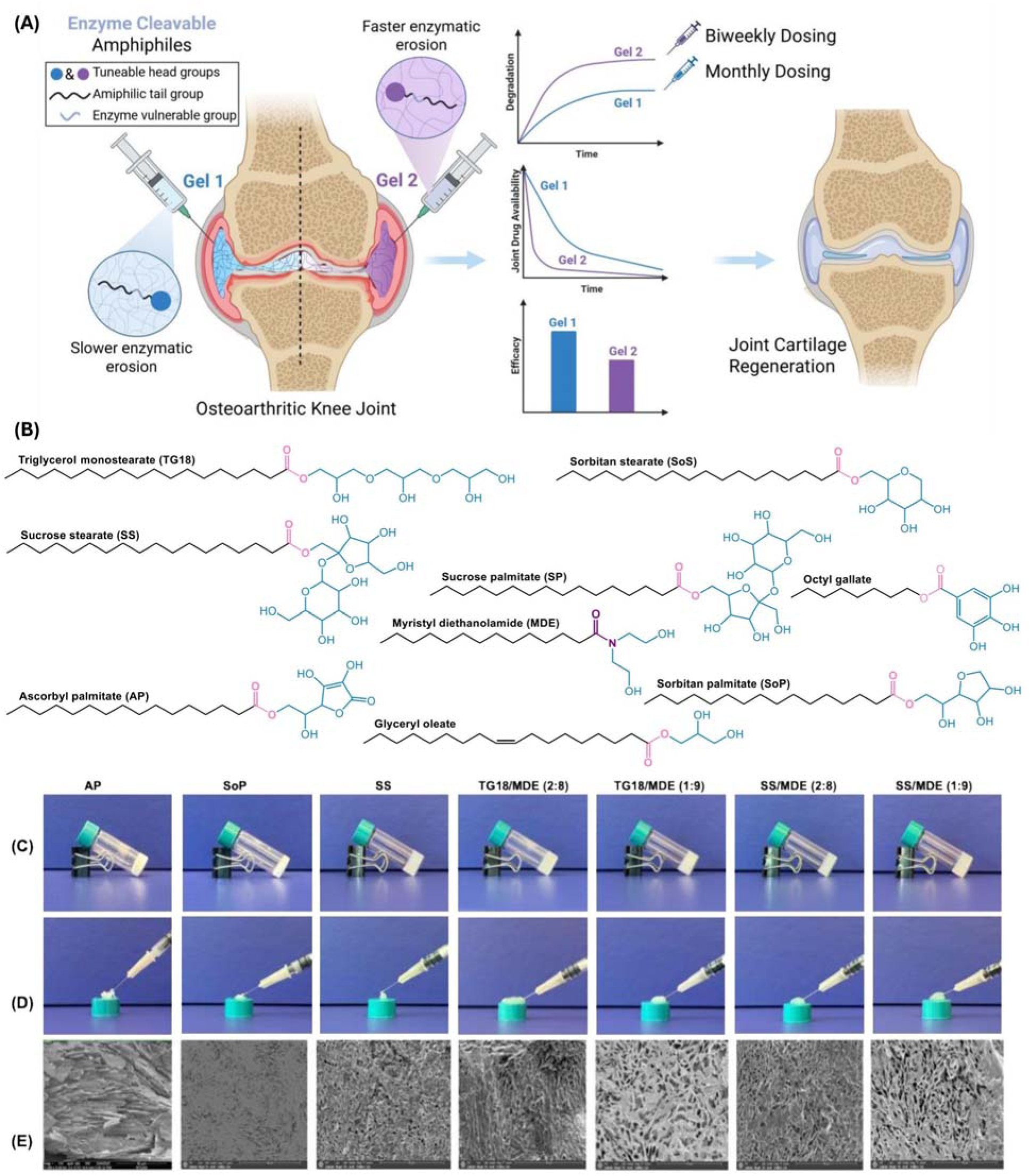
Design and characterization of tunable supramolecular hydrogels for intra-articular drug delivery. **(A)** Schematic illustration of the design concept: small-molecule amphiphiles comprising a hydrophilic head, hydrophobic tail, and enzyme-cleavable ester linkage self-assemble into supramolecular hydrogels capable of intra-articular injection. By modulating molecular composition, drug release kinetics can be tuned to enhance local retention and therapeutic efficacy. **(B)** Chemical structures of selected amphiphiles demonstrates variation in head group chemistry, hydrophobic tail length, and ester linkage susceptibility. **(C)** Vial tilting method to demonstrate hydrogel formation from seven formulations (10% w/v). **(D)** Photographs showing the injectability of amphiphile-based hydrogels (10% w/v). All seven gel formulations maintained self-supporting structures and could be smoothly extruded through a 27-gauge needle, confirming their suitability for intra-articular delivery**. (E)** Representative high resolution scanning electron microscopy (HR-SEM) images of hydrogel microstructures. Distinct nanofibrillar and porous architectures are observed across amphiphile compositions, reflecting differences in molecular packing and self-assembly behavior.

To tune the release kinetics of L-006235, we developed a rational screening approach to identify small-molecule amphiphiles capable of forming biocompatible low-molecular-weight hydrogels (LMWGs) similar to the TG-18 hydrogel.^12,15^ Beginning with a list of biocompatible amphiphiles that are generally regarded as safe by FDA, we prioritized amphiphiles containing enzyme-cleavable ester linkages and systematically varied head group chemistry and tail length to modulate enzymatic susceptibility and assembly behavior^12,16–18^ Through an initial gelation screen, three amphiphiles - ascorbic palmitate, sorbitan palmitate, and sucrose stearate - were able to form robust hydrogels. We then introduce a second amphiphile myristyl diethanolamide (MDE), which lacks an enzyme cleavage linkage, to mix with TG-18 or sucrose stearate at different ratios to form co-assembled hydrogels. In vitro release assays revealed that differential structures drove tunable drug release kinetics, with sucrose stearate (SS) hydrogels exhibiting the most sustained profile. In vivo imaging in healthy mice and mice with DMM surgery confirmed that SS gels doubled the intra-articular residence time compared to TG-18 hydrogels. When tested in the mouse model of post-traumatic osteoarthritis, SS/L-006235 hydrogels reduced cartilage degeneration scores by approximately 50% relative to vehicle and by ∼30% compared to TG-18/L-006235 formulations delivered at the same four-week dosing interval. Notably, the TG-18 hydrogel failed to confer same level of protection when administered every four weeks, whereas the SS hydrogel maintained therapeutic efficacy with monthly dosing. Together, these results establish a modular supramolecular hydrogel platform that enables clinically relevant tuning of intra-articular drug release kinetics. We hypothesized that local release kinetics, rather than residence time alone, would be a critical determinant of disease-modifying efficacy in PTOA.

## Results

### Screening LMWGs for self-assembly

The release of drugs that are non-covalently encapsulated in hydrogels primarily occurs through a combination of diffusion and material degradation. To precisely modulate the release kinetics of such systems, we sought to identify small-molecule amphiphiles featuring tunable structural parameters, including diverse hydrophilic head groups, varying hydrophobic tail lengths, and enzyme-cleavable ester linkages. Guided by these design principles, we systematically screened the biocompatible small molecules that are *Generally Recognized as Safe* (GRAS) and are commercially available in large quantities, we selected seven candidate amphiphiles in addition to triglycerol stearate (TG-18) that met our criteria (Figure 1B): ascorbyl palmitate (AP), glyceryl monooleate (GO), octyl gallate (OG), sorbitan palmitate (SM), sorbitan stearate (SoS), sucrose palmitate (SP), and sucrose stearate (SS).

To evaluate whether these small molecule amphiphiles enable the formation of hydrogels, each amphiphile was dissolved in dimethyl sulfoxide (DMSO, 10% w/v) at a temperature of 55-60°C to yield a clear solution, followed by addition of ultrapure water (90% v/v) and vortex mixing.^12,19^ Hydrophobic drugs such as L-006235 were incorporated during this dissolution process. The resulting mixtures were cooled to room temperature and assessed for gel formation; formulations that resisted flow upon vial inversion were classified as hydrogels. Among the seven candidates, AP, SM, and SS successfully formed self-supporting gels at 10% (w/v) (Figure 1C).

To further refine drug release kinetics, we adopted a co-assembly strategy in which primary amphiphiles (TG-18 or SS) were blended with a secondary amphiphile, myristyl diethanolamide (MDE), which lacks an enzyme-cleavable bond. By adjusting the MDE ratio (1:9 to 3:7, w/w), we also generated mixed TG-18/MDE and SS/MDE hydrogels (Figure 1C, Figure S1). All seven resulting hydrogel formulations exhibited smooth injectability through a 27-gauge needle (Figure 1D), underscoring their suitability for intra-articular administration in murine models.^20^ Scanning electron microscopy (SEM) revealed distinct nanofibrillar and porous network architectures among the various amphiphile systems (Figure 1E), reflecting differences in molecular packing and self-assembly behavior.

### Physical characterization of hydrogels

To evaluate the viscoelastic properties of seven distinct hydrogel formulations, we conducted strain sweep and dynamic frequency sweep tests using a rotational rheometer (Figure 2A). Strain sweeps from 0.01% to 50% were first performed at a constant frequency of 1 Hz to define the linear viscoelastic (LVE) region, the range within which the hydrogel structure remains intact and behaves predominantly elastically (G′ > G′′) (Figure 2B-H).^21^ The flow point, defined as the strain at which the material transitions from solid-like to liquid-like behavior (G′ < G′′), was identified for each formulation. SS gels exhibited a higher flow point than AP and SP gels. Incorporating MDE into TG-18 or SS significantly reduced both G′ and G′′ within the LVE region but increased the flow point relative to the undoped SS and TG-18 formulations. Subsequently, dynamic frequency sweeps were performed from 0.1 to 5 Hz at a fixed strain of 0.5%, which fell within the LVE region for all hydrogel samples (Figure 2I-O). Across all formulations, G′ consistently exceeded G′′, and both moduli increased with frequency, indicating frequency-dependent stiffening typical of viscoelastic gels.

**Figure 2.**
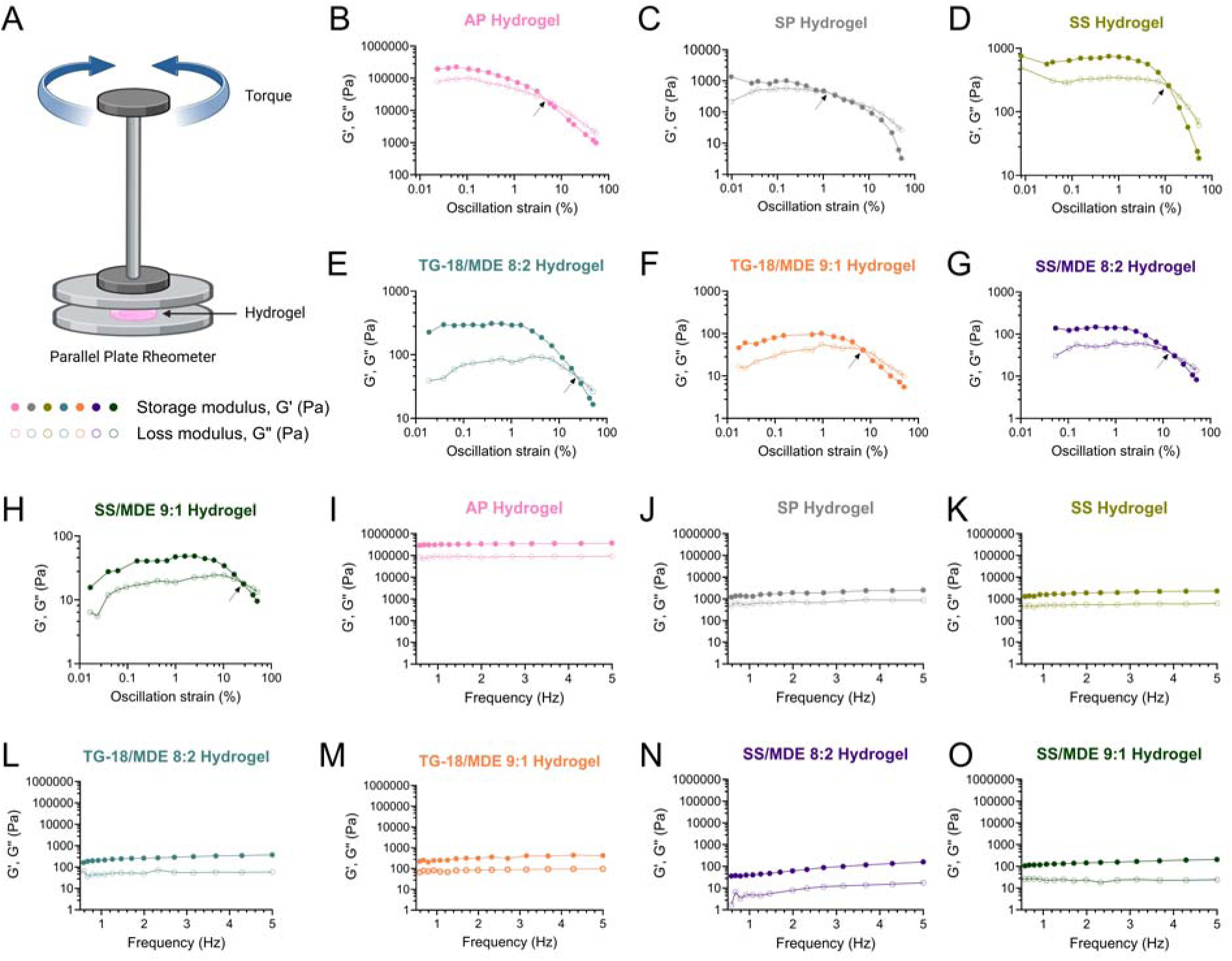
Mechanical property of self-assembled hydrogels. **(A)** Schematic illustration of physical characterization of hydrogels. **(B-H)** Storage modulus (G’) and loss modulus (G’’) of seven hydrogel formulations measured during a strain sweep performed at a fixed frequency of 1 Hz. **(I-O)** Storage modulus (G’) and loss modulus (G’’) measured during the dynamic frequency sweep of seven hydrogel formulations. Rheological measurements were performed to evaluate the viscoelastic behavior and mechanical stability of the self-assembled hydrogels. Data shown are representative of at least three independent experiments.

### In vitro and in vivo drug release kinetics

To evaluate the drug release kinetics of different hydrogel formulations, we loaded each hydrogel with L-006235, a small-molecule inhibitor of cathepsin-K, a key protease implicated in cartilage degradation as a model DMOAD. The hydrogels were prepared from different amphiphiles with L-006235 loaded at a concentration of 15 mg/mL. High-resolution scanning electron microscopy (HR-SEM) revealed higher-order assemblies within the drug-loaded hydrogels, confirming the formation of organized supramolecular networks (Figure S2). Drug release profiles were subsequently evaluated by incubating each hydrogel formulation in an esterase-rich environment containing T. lanuginosus lipase (200 U/mL) at 37 °C for 30 days, with enzyme replenishment every seven days (Figure 3A-G. The cumulative release data (Figure 3I-J) demonstrated distinct, formulation-dependent kinetics. AP and SP hydrogels exhibited rapid release, with over 75% of encapsulated drug released by day 30. In contrast, sucrose stearate (SS) hydrogels displayed the most sustained release, with ∼30% drug release by day 7 and a gradual increase to ∼40% by day 30. Incorporation of myristyl diethanolamide (MDE) into TG-18 hydrogels slowed release relative to pure TG-18, achieving ∼40% release at day 7 with extended release thereafter. MDE-doped SS hydrogels exhibited release kinetics similar to the undoped SS formulation over the same 30-day period (Figure 3J).

**Figure 3.**
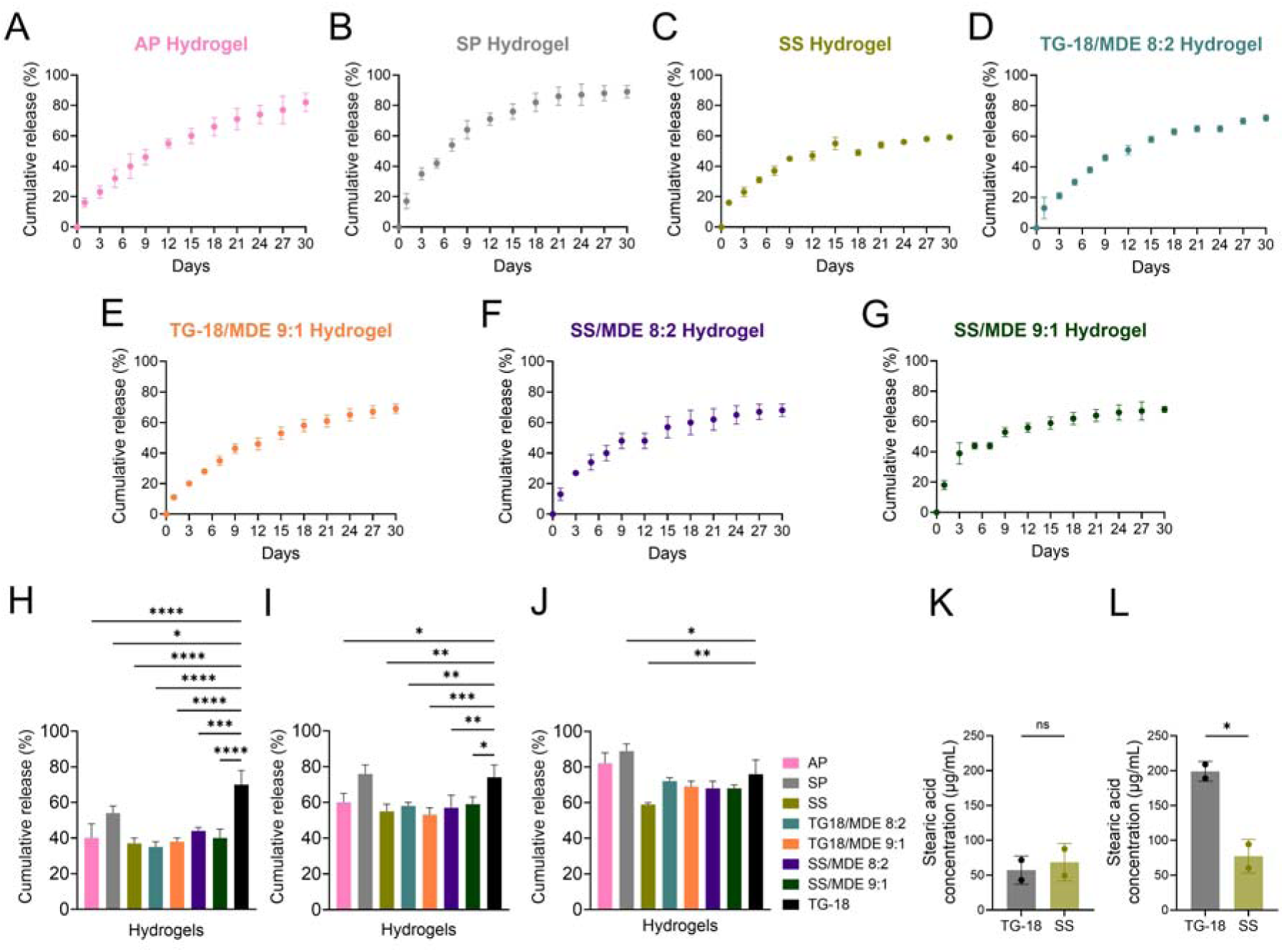
In vitro release kinetics of L-006235 from amphiphile-based hydrogel formulations. **(A)** ascorbic palmitate (AP), **(B)** sorbitan palmitate (SM), **(C)** sucrose stearate (SS), **(D)** TG-18/MDE (8:2 w/w), **(E)** TG-18/MDE (9:1 w/w), **(F)** SS/MDE (8:2), **(G)** SS/MDE (9:1). Hydrogels were incubated in PBS at 37 °C, and released drug was quantified at predetermined time points. **(I)** Percentage of cumulative drug release from all seven hydrogel formulations compared with TG-18 at day 7. **(H)** Percentage of cumulative drug release from all seven hydrogel formulations compared with TG-18 at day 15. **(J)** Percentage of cumulative drug release from all seven hydrogel formulations compared with TG-18 at day 30. **(K)** Stearic acid concentration of degraded TG-18 and SS hydrogels at day 0. **(L)** Stearic acid concentration of degraded TG-18 and SS hydrogels at day 6. Data represent mean ± SD of independent experiments.

Notably, TG-18 hydrogels exhibited faster drug release than SS hydrogels, which can be attributed to differences in their molecular structures and susceptibility to esterase-mediated degradation. TG-18 contains multiple glycerol–stearate ester linkages that are readily accessible to enzymatic hydrolysis, leading to faster network disassembly and accelerated drug release. In contrast, the sucrose-based architecture of SS forms a more stable supramolecular network in which ester bonds are less accessible, resulting in slower degradation and prolonged release. To directly assess enzymatic degradation, the release of stearic acid generated by ester bond cleavage was quantified by HPLC following incubation with esterase. After 6 days of incubation, TG-18 hydrogels released several-fold higher levels of stearic acid than SS hydrogels (Figure 3K-L), indicating substantially greater hydrolytic degradation. These results confirm that the faster drug release observed from TG-18 hydrogels is driven by their enhanced esterase-mediated degradation, whereas the slower degradation of SS hydrogels contributes to their more sustained release profile.

To evaluate the in vivo release and retention of encapsulated agents, we selected two hydrogel formulations, TG-18 and sucrose stearate (SS), that exhibited distinct release kinetics in vitro. Each hydrogel was co-loaded with the cathepsin-K inhibitor L-006235 (15 mg/mL) and a near-infrared fluorescent tracer, 1,1′-dioctadecyl-3,3,3′,3′-tetramethylindotricarbocyanine iodide (DiR, 1 mg/mL).^12^ A 4 µL aliquot of each hydrogel was injected intra-articularly (IA) into both knees of healthy (Figure 4A-D) and DMM (Figure 4E-G, Figure S3) C57BL/6 mice (day 0), and the fluorescence signal was monitored longitudinally using in vivo imaging (IVIS) for up to four weeks to assess hydrogel retention and clearance within the joint space. The SS hydrogel exhibited markedly prolonged intra-articular residence compared to TG-18, both in healthy and DMM mice, with detectable signal persisting up to 28 days post-injection, whereas fluorescence from TG-18 gels diminished by day 14, indicating approximately a twofold increase in residence time for SS.

**Figure 4.**
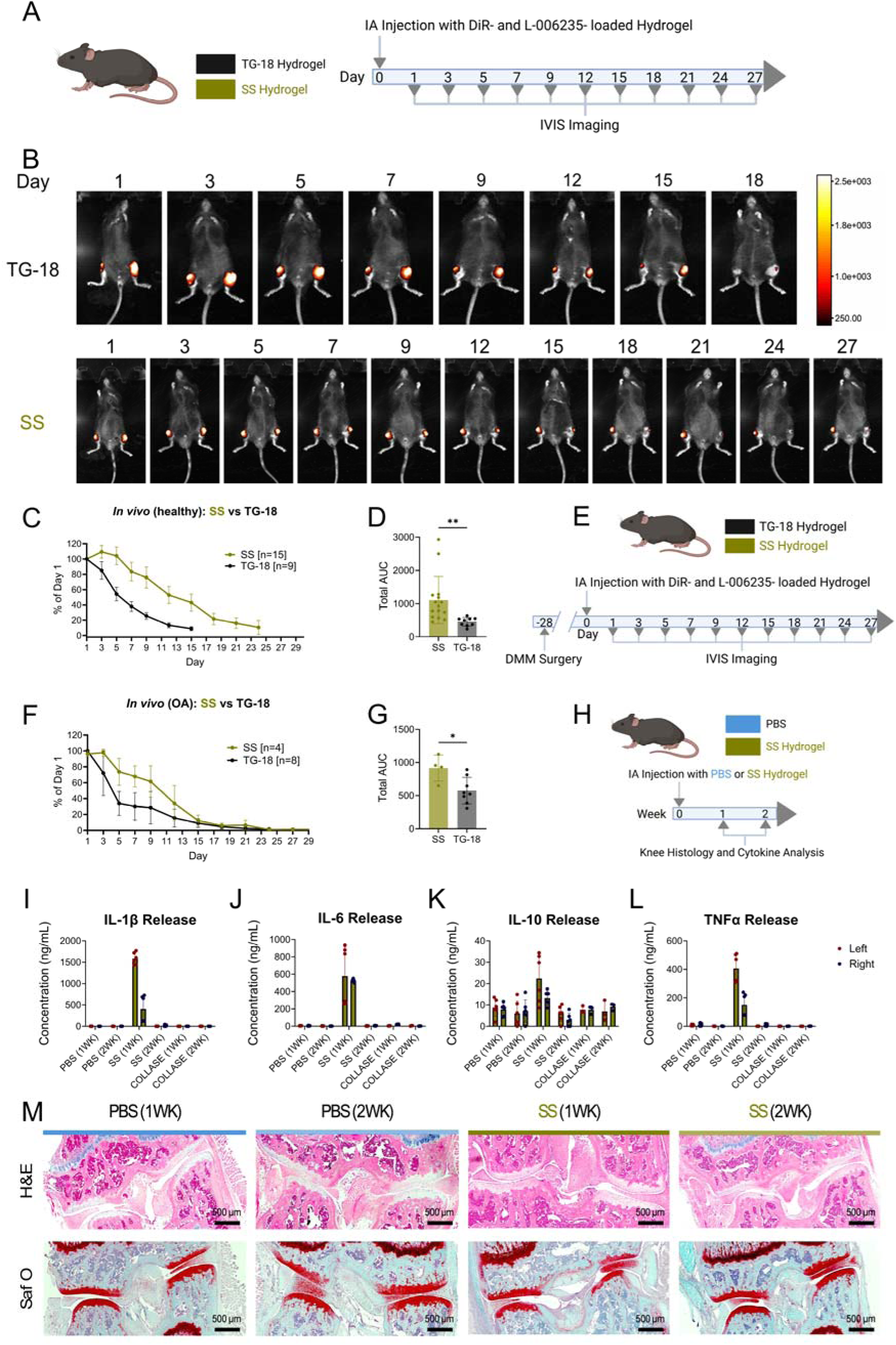
In vivo retention and biocompatibility of TG-18 and sucrose stearate (SS) hydrogels following intra-articular injection. **(A)** Experimental outline for longitudinal in vivo fluorescence imaging to evaluate retention of TG-18 and SS hydrogel following intra-articular injection into mouse knee joints. **(B)** Representative IVIS images showing fluorescence retention of TG-18 and SS hydrogels over time. **(C)** Quantitative analysis of normalized fluorescence intensity measured over injected knee joints over time. SS hydrogels maintained detectable fluorescence signal up to 28 days post-injection, doubling residence time relative to TG-18 hydrogels. Data are presented as mean ± SD (n = 10 joints). **(D)** Experimental timeline for evaluation of hydrogel biocompatibility following intra-articular injection of PBS or SS hydrogel. Joint tissues were collected at 1 and 2 weeks post-injection for histological analysis and cytokine quantification. **(E-L)** Quantification of inflammatory cytokines (IL-10, TNF-α, IL-1, IL-1β) in joint lysates collected at 1 and 2 weeks post-injection. A transient inflammatory response was observed at week 1 and resolved by week 2. **(M)** Representative histological images of joint sections stained with H&E and Safranin O/Fast Green demonstrating absence of synovial inflammation, cartilage degeneration, or tissue damage following SS hydrogel injection. Scale bar: 500 μm.

To assess the biocompatibility of the hydrogel formulations, we quantified inflammatory cytokines (IL-6, TNF-α, and IL-1β) and anti-inflammatory cytokine IL-10 at 1 and 2 weeks post-injection and performed histological evaluation of the joint tissues (Figure 4H-L). At one week, SS-injected joints showed a transient elevation in cytokine levels consistent with an acute inflammatory response typical of intra-articular administration. By two weeks, cytokine levels had returned to baseline, indicating that SS hydrogels were well-tolerated and did not elicit chronic inflammation. Histological analysis of knee sections revealed no evidence of synovial inflammation or tissue toxicity by hematoxylin and eosin (H&E) staining, and Safranin O staining confirmed preserved cartilage integrity in all groups (Figure 4M).^13,14^

### Efficacy of L-006235 gel to reduce PTOA progression

To evaluate the therapeutic outcomes of drug loaded hydrogels *in vivo*, we first validated the release kinetics of two gel formulations in a mouse model of PTOA. PTOA was induced *via* destabilization of the medial meniscus (DMM) surgery.^22^ Four weeks after the surgery, we injected TG-18 and SS (each loaded 15mg/mL L-006235, 1 mg/mL DiR) to the surgical treated knee joint and monitor the drug retention over time. SS still showed more sustained release than TG-18, with twice longer half-lives (Figure S3). Then to evaluate how local drug release kinetics in the joint impact therapeutic outcomes *in vivo,* we tested drug loaded TG-18 and sucrose stearate hydrogels with the same dosing scheme in a post-traumatic osteoarthritis (PTOA) mouse model. Treatments were initiated at week 5 post-surgery (early OA phase) (Figure 5A). The following treatment groups were established (all injections intra-articularly in the affected knee joint): 1) Sham: Mice underwent sham surgery (no DMM) and received no drug treatment, 2) DMSO biweekly injections of vehicle (DMSO/water), 3) TG-18 hydrogel loaded with L-006235, injected every 2 weeks (frequent dosing), 4) TG-18 hydrogel with L-006235, injected every 4 weeks, 5) SS hydrogel loaded with L-006235, injected every 4 weeks, 6) injection of SS hydrogel without drug (every 4 weeks). All mice were sacrificed at 12 weeks post-surgery (8 weeks after the first injection). Prior to histology, joints were assessed by micro-CT to examine structural changes in bone. Knees were then decalcified, sectioned, and stained with Safranin O/Fast Green to visualize cartilage proteoglycan content. Cartilage degeneration was scored in a blinded fashion using the Osteoarthritis Research Society International (OARSI) histopathology scoring system, which grades the severity of cartilage lesions. ^23^

**Figure 5.**
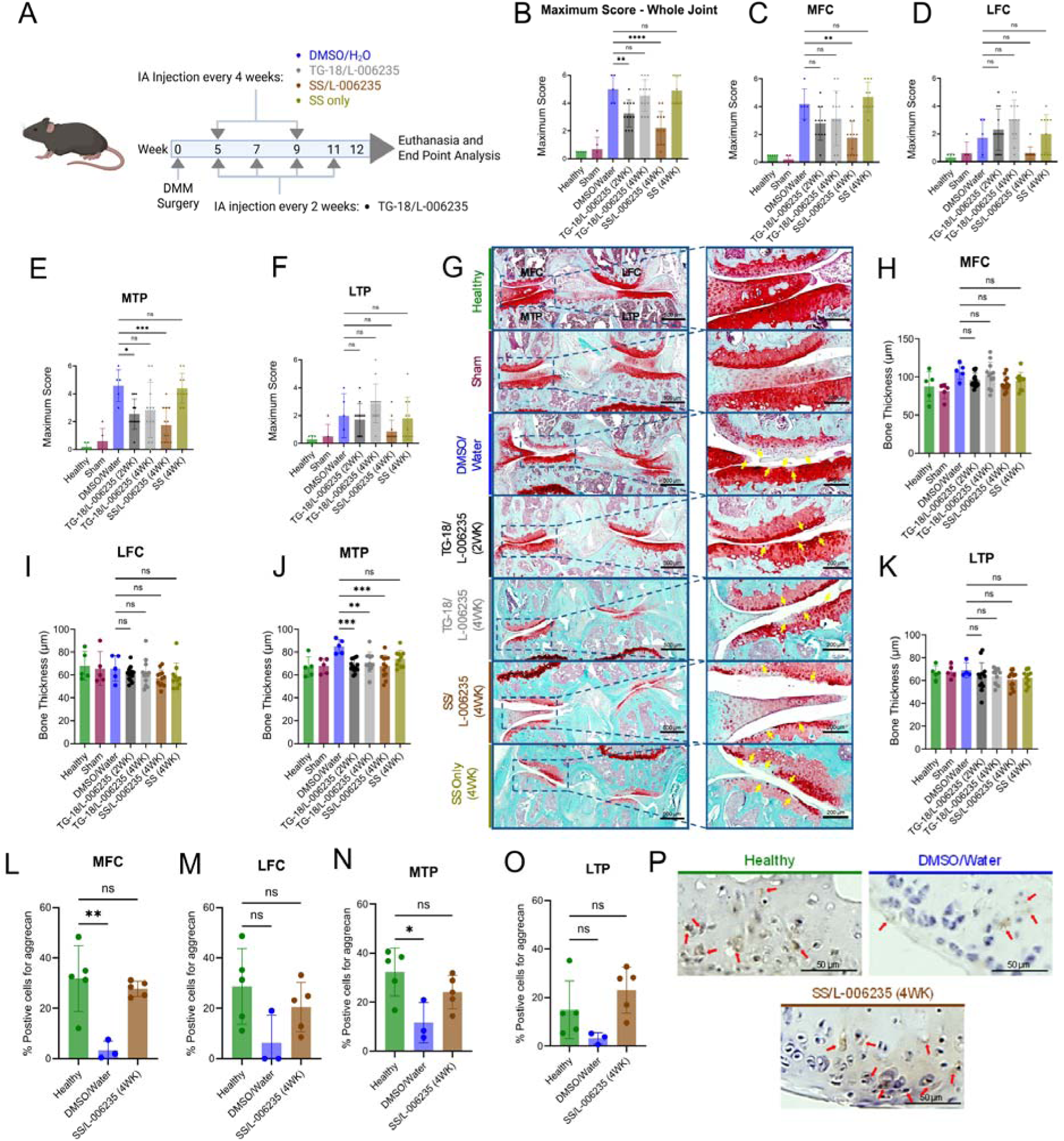
Therapeutic efficacy of L-006235-loaded hydrogels in a post-traumatic osteoarthritis (PTOA) mouse model. **(A)** Experimental outline for evaluation of therapeutic efficacy in mice with surgically induced PTOA. Mice underwent destabilization of the medial meniscus (DMM) surgery followed by intra-articular administration of treatment formulations and longitudinal assessment of disease progression. **(B–F)** Quantification of Osteoarthritis Research Society International (OARSI) histopathology scores for the entire joint and individual joint compartments, including the medial femoral condyle (MFC), lateral femoral condyle (LFC), medial tibial plateau (MTP), and lateral tibial plateau (LTP). **(G)** Representative Safranin O/Fast Green-stained histological sections from each treatment group demonstrating differences in cartilage preservation and degeneration severity. Arrows indicate regions of cartilage erosion and structural damage. Scale bars: 500 µm in low-magnification images and 200 µm in zoomed-in images. **(H–K)** Quantitative analysis of subchondral bone thickness in each joint quadrant measured from micro-computed tomography (µCT) images. **(L-O)** Quantitative analysis of aggrecan positive cells from each treatment group, including MFC, LFC, MTP, and LTP. **(P)** Representative immunofluorescence staining for aggrecan in joint sections from each treatment group. Arrows denote positive staining. Scale bar: 20 µm. P-values for OARSI scoring were determined using the Kruskal–Wallis nonparametric test, while subchondral bone thickness measurements were analyzed using one-way ANOVA with Tukey’s post hoc analysis following confirmation of normality. **P < 0.01 and ***P < 0.001.

Histological analysis revealed a robust chondroprotective effect of the L-006235-loaded hydrogels. Mice treated with the SS/L-006235 hydrogel (dosed every four weeks) exhibited a significantly lower cartilage degeneration score compared to DMSO group, the maximum OARSI score for the entire joint was reduced by ∼56% in the SS/L-006235 group relative to the DMSO control (p < 0.01) (Figure 5B). This indicates that the sustained-release SS hydrogel markedly preserved cartilage structure in vivo, whereas untreated PTOA joints developed severe degeneration. Notably, the SS/L-006235 group also showed significantly better cartilage protection than the TG-18/L-006235 group that was injected every four weeks, suggesting that the SS hydrogel’s sustained release profile provided superior therapeutic efficacy (Figure 5B). The empty SS hydrogel (no drug) did not significantly improve scores over vehicle, confirming that the benefit was due to L-006235 release rather than the material itself. The therapeutic outcomes differed depending on dosing frequency for the TG-18 hydrogel. Mice receiving TG-18/L-006235 every 2 weeks showed a greater reduction in OARSI scores than those receiving TG-18/L-006235 every 4 weeks (Figure 5B).

Consistent with the DMM model’s pathology, DMSO/H2O-treated PTOA mice showed more severe cartilage loss in the medial knee compartments than in the lateral ones (Figure 5C-G). OARSI scores in the DMSO control group were markedly higher in the medial quadrants (medial femoral condyle (MFC) and tibial plateau (MTP)) than in the lateral quadrants, reflecting concentrated cartilage degeneration medially. Treatment with SS/L-006235 hydrogel preferentially protected these vulnerable medial regions. The SS/L-006235 (4w) group had significantly lower cartilage degeneration scores in the medial compartments compared to the DMSO group (p < 0.05), indicating that much of the medial cartilage was preserved. In the lateral compartments, where degeneration was milder, the SS/L-006235 treatment did not show a statistically significant difference from controls – suggesting the therapy primarily mitigated the more pronounced medial damage. The TG-18/L-006235 groups showed a similar trend of medial protection, with the biweekly TG-18 treatment yielding the better medial improvement (consistent with its overall higher efficacy when dosed frequently). We also assessed subchondral bone remodeling by measuring subchondral bone plate (SBP) thickness in each quadrant of the knee joint (Figure 5H–K). Consistent with previous reports of osteoarthritis-associated subchondral bone sclerosis, DMM mice treated with DMSO/water exhibited significantly increased SBP thickness in the medial tibial plateau (MTP) compared with sham-operated controls, whereas no significant differences were observed in the medial femoral condyle (MFC), lateral femoral condyle (LFC), or lateral tibial plateau (LTP) regions.^12,24^ The increase in MTP SBP thickness is consistent with the more severe cartilage degeneration observed histologically in the medial compartment of the joint. Treatment with TG-18/L-006235 administered either every 2 weeks or every 4 weeks significantly reduced SBP thickness in the MTP relative to the DMSO/water group, indicating attenuation of pathological subchondral bone remodeling. In contrast, neither SS/L-006235 nor blank SS hydrogel significantly altered SBP thickness despite showing a modest trend toward improvement. These findings suggest that sustained delivery of L-006235 from TG-18 hydrogels effectively mitigates OA-associated changes in the subchondral bone, complementing the observed improvements in cartilage preservation and overall joint pathology.

To further assess the effects of L-006235-loaded SS hydrogels on cartilage matrix components, we performed immunofluorescence staining for aggrecan key markers of cartilage degradation and repair. In the DMSO-treated PTOA group, we observed a much higher reduction in aggrecan levels, consistent with the extensive cartilage degradation typically seen in the medial compartments of PTOA joints (Figure 5L-P). Treatment with SS/L-006235 hydrogel significantly restored aggrecan expression. Overall, the sustained-release SS hydrogel loaded with L-006235 demonstrated superior in vivo efficacy in the PTOA model, significantly reducing cartilage degeneration and preserving matrix content with only monthly injections. By contrast, the faster release gel formulation TG-18 hydrogel required more frequent dosing to achieve comparable benefits, and even then, the SS hydrogel-treated mice showed the lowest cartilage lesion scores.

## Discussion

To our knowledge, this is the first study to systematically evaluate how local drug release kinetics influence disease-modifying outcomes in PTOA. Our findings demonstrate that engineering release kinetics, rather than simply extending intra-articular residence time, can materially improve therapeutic efficacy. In this study, we explored the impact of local drug release kinetics on therapeutic efficacy in a mouse model of post-traumatic osteoarthritis (PTOA). By comparing different hydrogel formulations loaded with L-006235, a small molecule inhibitor of cathepsin-K, we demonstrated that the sustained-release SS hydrogel provided superior therapeutic efficacy with minimal dosing, while the more frequent dosing regimen of TG-18 hydrogels offered less prolonged joint retention and less cartilage protection. These findings highlight the importance of optimizing local drug release kinetics for the treatment of joint diseases like PTOA, where prolonged drug exposure is essential for maintaining cartilage integrity.

Prior studies have shown that hydrogels and nanoparticle-based delivery systems can prolong the residence time of therapeutics within the joint, offering an advantage over conventional treatments that typically require frequent injections. However, how local release kinetics in the joint affect the therapeutic efficacy is not understood. Our results support the notion that drug release kinetics play a crucial role in the therapeutic outcomes of intra-articular drug delivery systems.

Importantly, this work provides one of the clearest demonstrations to date that rational control of intra-articular release kinetics can directly influence disease progression and therapeutic outcomes in osteoarthritis. Through systematic screening of the biocompatible and commercially available small molecule amphiphiles for the development of hydrogel formulations, we identified seven hydrogel formulations that were both biocompatible and capable of providing controlled drug release. These small molecule amphiphiles are not only safe for therapeutic use but also have strong translational potential. This rigorous screening process helped us identify the sucrose stearate (SS) hydrogel formulation, which demonstrated superior sustained drug release and longer retention in the joint, making it an ideal candidate for disease-modifying osteoarthritis therapy.

By comparing two hydrogel formulations with distinct release kinetics (TG-18 and SS) in vivo, we were able to directly assess how sustained drug release influences therapeutic efficacy. The use of micro-CT imaging and Safranin O staining provided detailed insights into joint structure and cartilage preservation, making it possible to quantify the therapeutic effects of L-006235-loaded hydrogels. Additionally, the use of a clinically relevant PTOA model and a well-validated histological scoring system (OARSI) ensures that our findings have strong translational potential for future clinical applications.

Moreover, the study design included a comprehensive range of treatment regimens, including dosing every two weeks, four weeks, and a hydrogel-only control group. This robust comparison offers a nuanced understanding of how drug release kinetics directly affect cartilage preservation, providing critical data for future therapeutic strategies in PTOA and potentially other degenerative joint diseases.

While our study provides compelling evidence for the potential of sustained-release hydrogels in PTOA treatment, there are certain limitations. First, the DMM-induced PTOA model is a well-established and widely used model for studying osteoarthritis; however, its pathophysiology may not capture the full range of inflammatory and degenerative processes seen in humans. This limits the generalizability of our findings to clinical populations, and future studies using more diverse models that mimic human OA progression are needed to further validate these results. Another limitation is the single dose of L-006235 used in our study. While this dose was selected based on previous studies of L-006235’s efficacy in reducing cartilage degradation, dose optimization and dose escalation studies are necessary to determine the ideal concentration for long-term therapeutic benefit. Additionally, this study focused on small molecule inhibitors for PTOA, future research exploring alternative drug candidates, such as biologics or RNA-based therapies, within these hydrogel formulations could provide new avenues for disease-modifying therapies.

This study establishes local release kinetics as a critical determinant of disease-modifying efficacy in post-traumatic osteoarthritis and identifies release kinetics as a key design parameter for intra-articular therapeutics. Using a modular supramolecular hydrogel platform composed of biocompatible GRAS amphiphiles, we demonstrate that rationally engineered release profiles can substantially improve cartilage preservation while reducing dosing frequency. Notably, sustained release from the optimized formulation enabled effective monthly administration while improving cartilage preservation, addressing a major challenge in intra-articular therapy. Collectively, these findings provide a framework for the development of long-acting disease-modifying therapies for osteoarthritis and other chronic joint diseases.

## Methods

### Hydrogel formulation

Low–molecular-weight amphiphile hydrogels were prepared as described previously, with minor modifications. To prepare 1 mL of hydrogel, amphiphile (100 mg) was weighed into a glass scintillation vial, followed by the addition of dimethyl sulfoxide (DMSO, 200 µL).^12^ The mixture was heated to 55–60 °C until complete dissolution was achieved and a clear solution was obtained.

While the solution was still hot, ultrapure water (800 µL) was added immediately. The mixture was vortexed for 5 s, briefly reheated for an additional 5 s, and vortexed again to obtain a homogeneous solution. The vial was then placed on a flat surface and allowed to cool at room temperature for 15–30 min, resulting in hydrogel formation. Gelation was confirmed by the absence of gravitational flow upon inversion of the vial.

For L-006235 gel or L-006235/DiR co-loaded hydrogel, L-006235 (Tocris Bioscience), with or without 1,1′-dioctadecyl-3,3,3′,3′-tetramethylindotricarbocyanine iodide (DiR) (Thermo Fisher Scientific), was added together with the amphiphile to achieve a final concentration of 10–20 mg/mL L-006235 (1–2% w/v) or 100 μg/mL DiR (0.01% w/v).

### In vitro release kinetics

Drug release was evaluated using a dialysis bag method as previously reported. Briefly, L-006235–loaded gel (50 μL, 15 mg/mL) was transferred into dialysis tubing with an 8–10 kDa molecular weight cut-off (Spectrum Labs) and initially suspended in 550 μL of PBS. To mimic the joint environment, 200 μL of either 100% human osteoarthritic (OA) synovial fluid (Articular Engineering) or an equivalent volume of PBS was added to the dialysis bag at designated time points. The sealed dialysis bags containing the hydrogel and release medium were then immersed in 40 mL of PBS as the sink medium and incubated at 37 °C with orbital shaking at 150 rpm.

At each sampling time point, 1 mL of the sink medium was collected and immediately replaced with an equal volume of fresh PBS to maintain sink conditions. Collected samples were lyophilized, reconstituted in 250 μL of DMSO, and analyzed by high-performance liquid chromatography (HPLC) using an Agilent 1260 Infinity II system equipped with a Zorbax SB-C18 column (250 × 4.6 mm, 5 μm).

The cumulative percentage of drug released at time t (Ct) was calculated by accounting for the amount of drug present in the sink medium at time t (At) and the cumulative amount removed in aliquots collected at previous time points (Ai). The total drug loading was 750 μg per 50 μL of L-006235 gel, as confirmed by HPLC.

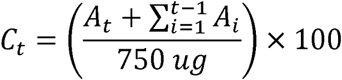

*where:*

- Ct represents the cumulative percentage of drug released at time t,
- At denotes the amount of drug present in the sink medium at time t,
- Ai corresponds to the amount of drug removed in each 1 mL aliquot collected at earlier time points i, and
- 750 μg is the total drug content in 50 μL of the L-006235 gel, as confirmed by HPLC analysis.

### Esterase-mediated degradation of TG-18 and SS hydrogels

Two replicates of 50 μL of TG-18 and SS hydrogels were cured at 37 ℃ for 45 min. To each hydrogel, 2 mL of 200 U/mL of porcine liver esterase in PBS were added. After 6 days, each hydrogel was broken using a pipette, centrifuged at 5000 rpm for 5 min. The supernatant was discarded, and replaced with 2 mL of ethanol. The pellet in ethanol was shaken at 37 ℃ for 2 days, broken using a pipette, and filtered through a 0.22 μm PTFE syringe filter. As a control, the process was repeated using PBS instead of the esterase solution.

The ethanol solutions were analyzed using Agilent 6530 QTOF LCMS, with a C18 RP LC column. A gradient elution was performed using a combination of 10 mM ammonium formate in water and 10 mM ammonium formate in acetonitrile. Stearic acid was quantified by selecting for its [M-H]*-* ion, with an m/z value of 283.2637.

### Rheological characterization

Rheological measurements were performed using a rotational rheometer (Discovery HR-2, TA Instruments) equipped with a cone–plate geometry (parallel plate, 15 mm diameter), as described previously with minor modifications. All measurements were conducted at 37 °C to mimic physiological conditions. Approximately 500 µL of freshly prepared hydrogel was loaded onto the rheometer plate for each measurement.

Strain sweep experiments were performed to determine the linear viscoelastic (LVE) region of each hydrogel formulation. Strain was varied from 0.01% to 50% at a constant frequency of 1 Hz, and the storage modulus (G′) and loss modulus (G″) were recorded. The LVE region was defined as the strain range over which G′ exceeded G″ and remained independent of strain. The flow point was identified as the strain at which G′ crossed G″, indicating a transition from solid-like to liquid-like behavior.^25,26^

Dynamic frequency sweep measurements were subsequently performed within the LVE region at a constant strain of 0.5 %. Frequency was varied from 0.1 to 5 Hz, and G′ and G″ were recorded to evaluate the frequency-dependent viscoelastic behavior of the hydrogels.

All rheological measurements were performed on freshly prepared hydrogels, and representative data were obtained from at least three independent samples per formulation.

### Animals

All animal experiments were performed using 6–10-week-old male C57BL/6J mice (Jackson Laboratory). Mice were housed under standard 12 h light–dark cycles with ad libitum access to food and water. All procedures were approved by the Institutional Animal Care and Use Committee (IACUC) at Brigham and Women’s Hospital and conducted in accordance with NIH and AAALAC guidelines.

### Scanning Electron Microscopy (SEM)

The microstructure of hydrogel formulations was characterized using scanning electron microscopy (SEM). For blank hydrogels, samples were frozen at −80 °C and lyophilized to remove water prior to imaging. The freeze-dried hydrogels were fractured to expose internal cross-sections, mounted on aluminum stubs using carbon tape, and sputter-coated with gold for 60 s. Imaging was performed using an FEI Quanta 450 field-emission scanning electron microscope (FESEM) to evaluate pore architecture and surface morphology.

### Cryogenic Scanning Electron Microscopy (Cryo-SEM)

To preserve the native hydrated morphology of drug-loaded hydrogels and minimize artifacts associated with lyophilization, cryogenic SEM (cryo-SEM) was employed. Cryo-SEM imaging was conducted using a Thermo Scientific Quanta FEG SEM equipped with a Quorum PP3010T cryo-preparation and cryo-transfer system. A small amount of freshly prepared hydrogel was placed onto a specimen holder coated with colloidal graphite conductive paste, along with a minimal volume of OCT compound to enhance sample adhesion. Samples were mounted on the PP3010T Prepdek® workstation, plunge-frozen in liquid nitrogen, and transferred under vacuum to the cryo-preparation chamber.

Within the chamber, samples were gently fractured using a handle-mounted blade to expose internal structures, followed by controlled sublimation to remove surface ice using the system’s automated five-step protocol. The fractured surfaces were subsequently sputter-coated and transferred directly to the SEM chamber for imaging under cryogenic conditions. SEM and cryo-SEM images were acquired at low accelerating voltages to preserve fine structural features. Representative images were collected from multiple regions of each sample to ensure reproducibility of the observed nanofibrillar network morphology.

### In vivo biocompatibility

Local biocompatibility of the hydrogel formulations was evaluated in healthy mice following a single intra-articular (IA) injection. Mice received 4 µL of phosphate-buffered saline (PBS), sucrose stearate hydrogel, or collagenase-treated sucrose stearate hydrogel (+ control) injected into the knee joint.^27,28^ Animals were euthanized at 7 or 14 days post-injection for cytokine analysis and histological assessment.^23,29^

For cytokine analysis, synovial fluid was collected by lavage. Briefly, the joint space was washed eight times with 2.5 µL PBS to recover synovial fluid, and the pooled lavage was used for downstream analysis. Levels of inflammatory and anti-inflammatory cytokines (including IL-6, TNF-α, IL-1β, and IL-10) were quantified using multiplex electrochemiluminescence immunoassays (Meso Scale Discovery, MSD). For cytokine studies, three mice were used per treatment group per time point, resulting in a total of 18 mice.

For histological evaluation, knee joints were harvested at the same time points, fixed, decalcified, embedded in paraffin, and sectioned. Sections were stained with hematoxylin and eosin (H&E) and Safranin O/Fast Green to assess synovial inflammation, cartilage integrity, and overall joint morphology. Histological analysis was performed on 18 joints (18 mice) in a blinded manner.

### Destabilization of the Medial Meniscus (DMM) Model

The destabilization of the medial meniscus (DMM) model was established as previously described.^22^ Briefly, under anesthesia, the right knee joint was surgically exposed through a medial parapatellar incision adjacent to the patellar ligament, and the medial meniscotibial ligament (MMTL) was transected to induce joint destabilization. For sham-operated controls, the right knee joint was similarly exposed, but the MMTL was left intact.

### In vivo assessment of joint retention

For joint retention studies, TG-18 or SS hydrogels co-loaded with DiR were injected intra-articularly (4 µL) into mouse knee joints of healthy mice and DMM model.^30^ Fluorescence signals were monitored longitudinally using an in vivo imaging system (IVIS) at predefined time points for up to 28 days. Regions of interest were quantified, and signal intensities were normalized to day-0 values to assess hydrogel residence time within the joint.

### In vivo therapeutic efficacy study

Therapeutic studies were initiated 5 weeks post-DMM surgery (early OA phase). Mice received intra-articular injections according to the assigned treatment group: vehicle, TG-18/L-006235 (biweekly or monthly), SS/L-006235 (monthly), or empty hydrogel. All injections were administered at a volume of 4 µL into the operated knee. Animals were euthanized at 12 weeks post-surgery for joint analysis.

### Immunohistochemistry

Immunohistochemical staining was performed on joint tissue sections using an automated Leica Bond RX platform by a commercial histology service provider (iHisto, LLC). Slides were stained per marker on a per-slide basis using antibodies against aggrecan and matrix metalloproteinase-13 (MMP-13) following standardized protocols. Brightfield images were acquired using a Zeiss AxioScan 7 whole-slide scanner. Quantitative analysis of staining intensity was performed using ImageJ software.

### Histology

Following euthanasia, knee joints were harvested and placed into labeled histology cassettes using alcohol-resistant ink. Joints were dissected and fixed in 10% neutral buffered formalin for 24–48 h. Samples were then rinsed thoroughly with tap water and transferred to 70% ethanol for storage prior to imaging. Fixed samples were submitted for micro–computed tomography (μCT) scanning.

After completion of μCT imaging, samples were rinsed with tap water and transferred to formic acid–based decalcification medium. Decalcification was carried out for 2–3 days under gentle agitation on a shaker to ensure uniform decalcification while preserving cassette labeling. The decalcification solution was then replaced with fresh medium, and samples were decalcified for an additional 2–3 days. Upon completion, samples were rinsed thoroughly with tap water and transferred back to 70% ethanol.

Decalcified joints were subsequently submitted for paraffin embedding, sectioning, and histological processing. Sections were cut in the frontal plane and used for downstream histological and immunohistochemical analyses.

### Micro–Computed Tomography (μCT) Imaging

Following euthanasia, injected knee joints were harvested and fixed in 4% (w/v) paraformaldehyde for 24 h, followed by storage in 70% (v/v) ethanol. Samples were scanned using a μCT 35 system (Scanco Medical) equipped with μCT V6.1 software. Scans were acquired at an isotropic voxel size of 12 μm, an X-ray tube voltage of 55 kV, a current of 145 μA, and an integration time of 600 ms. Image reconstruction and analysis were performed using the manufacturer’s built-in software.

Three-dimensional reconstructions of knee joints were generated using a global bone mineral density threshold of 380 mg HA/cm³ and digitally sectioned along the frontal plane. Load-bearing regions of the joint were selected as regions of interest. Subchondral bone plate thickness was assessed by contouring the cortical bone of the medial tibial plateau while excluding calcified articular cartilage and osteophyte regions.^24^ Measurements were obtained from the medial and lateral regions of the tibial plateau and femoral condyle at the mid-facet section of the knee joint using Scanco Medical μCT V6.1 software.^22^

### Software and Statistical Analysis

IVIS images were captured and quantified using Bruker MI SE software version 7.5.2.22464. Histological slides were accessed using Zeiss Zen 3.13. Quantification of aggrecan immunohistological staining was performed using Fiji ImageJ version 1.54p. Data was graphed using GraphPad Prism version 10.6.1. Molecular structures of amphiphiles were obtained using ChemDraw version 17.0. Figure schematics were created using BioRender. ChatGPT Plus was used for formatting.

All statistical analyses and graphical representations were performed using GraphPad Prism. Comparisons between two experimental groups were conducted using a two-tailed Student’s t-test. For comparisons involving more than two groups, one-way analysis of variance (ANOVA) followed by Tukey’s post hoc test was applied.

For in vivo fluorescence imaging studies of L-006235/DiR co-loaded hydrogels, fluorescence signal decay was quantified by calculating the area under the curve (AUC) for each individual mouse, and group means were compared. In vitro drug release from the L-006235 hydrogel in phosphate-buffered saline (PBS) versus osteoarthritic (OA) synovial fluid was analyzed using two-way ANOVA, with time and sink condition (PBS or OA synovial fluid) as independent variables.

Normality of in vivo efficacy data was assessed using the Kolmogorov–Smirnov test. Because cartilage degeneration scores in some groups did not follow a normal distribution, statistical significance was evaluated using the Kruskal–Wallis test. Subchondral bone plate (SBP) thickness data were normally distributed and therefore analyzed using one-way ANOVA with Tukey’s post hoc test.

For analyses of cartilage degeneration scores and SBP thickness, the mean of each treatment group was compared with the mean of the DMSO/PBS-treated negative control group. A P value < 0.05 was considered statistically significant.

## Acknowledgments

NIH grant R01AR077718 (N.J.) Football Players Health Study grant (N.J., J.M.K., and J.E.). The Football Players Health Study is funded by a grant from the National Football League Players Association. The content is solely the responsibility of the authors and does not necessarily represent the official views of Harvard Medical School, Harvard University or its affiliated academic health care centers, the National Football League Players Association, or the Brigham and Women’s Hospital.

