## Supplement for "Systematic Engineering of Intra-Articular Drug Release Profiles Reveals a Key Determinant of Disease-Modifying Efficacy in Post-Traumatic Osteoarthritis"

^3^Harvard Medical School, Boston, MA 02115, USA.

^4^Broad Institute of MIT and Harvard, Cambridge, MA 02142, USA.

^5^Harvard–Massachusetts Institute of Technology Division of Health Sciences and Technology, Massachusetts Institute of Technology, Cambridge, MA 02139, USA

^6^Division of Engineering, New York University Abu Dhabi (NYUAD), Abu 7 Dhabi, UAE

^7^Department of Mechanical and Aerospace Engineering, Tandon School of Engineering, New York University, Brooklyn, NY 11201, USA

^7^Harvard Stem Cell Institute, Cambridge, MA 02138, USA.

^8^Division of Rheumatology, Inflammation and Immunity, Department of Medicine, Brigham and Women’s Hospital, Boston, MA 02115, USA.

*Corresponding authors. (J.G.); (N.B); (J.M.K.); (J.E.); (N.J.)

†These authors contributed equally to this work.

**This PDF file includes:**

Figs. S1 to S4

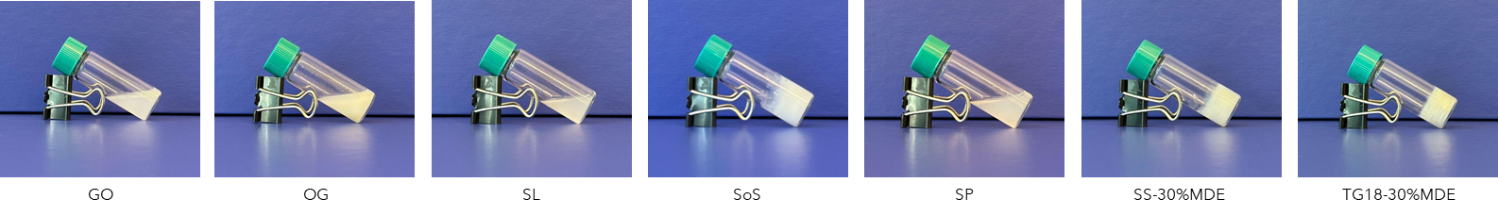

**Figure S1.** Photographic images illustrating the vial inversion (tube-tilt) test used to assess gelation behavior. Upon inversion, the sample (10% w/v) remains immobilized against gravity, indicating successful gel formation the sample remains flowable and continues to move/settle under gravity over time, indicating the formulation did not form a self-supporting gel.

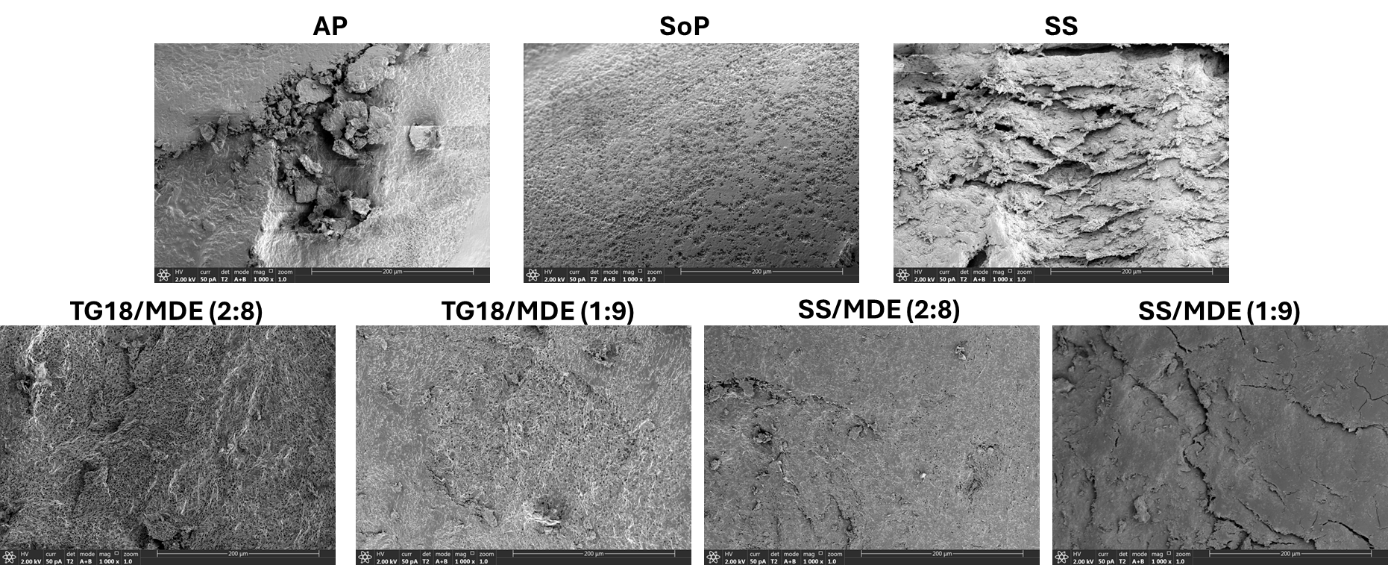

**Figure S2.** Representative high resolution scanning electron microscopy (HR-SEM) images of L-006235 loaded hydrogel microstructures. Distinct nanofibrillar and porous architectures are observed across amphiphile compositions, reflecting differences in molecular packing and self-assembly behavior.

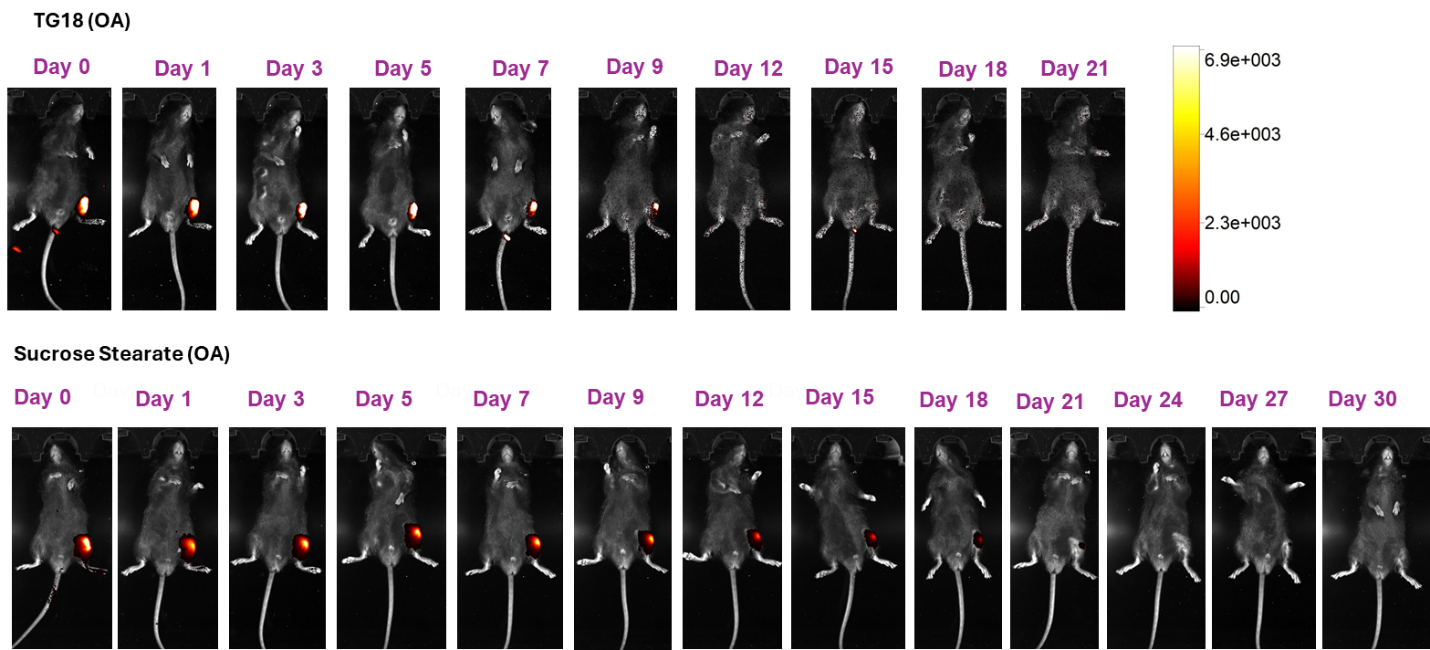

**Figure S3.** Representative longitudinal IVIS images following intra-articular injection of DiR-loaded sucrose stearate (SS) and TG-18 hydrogels into osteoarthritic (OA) joints.

**Figure S4.** Representative 2D μCT images of knee joints across experimental groups.
Images show the medial and lateral compartments of the tibial plateau and femoral condyles in healthy, sham, DMSO, TG-18 + L-006235 (dosed every 2 or 4 weeks), and sucrose stearate–based treatment groups. Subchondral bone plate thickness was quantified by contouring the cortical bone of the medial and lateral tibial plateau and femoral condyles while excluding calcified articular cartilage and osteophyte regions. Measurements were performed on the central slice of each contoured region using Scanco Medical μCT software (v6.1).

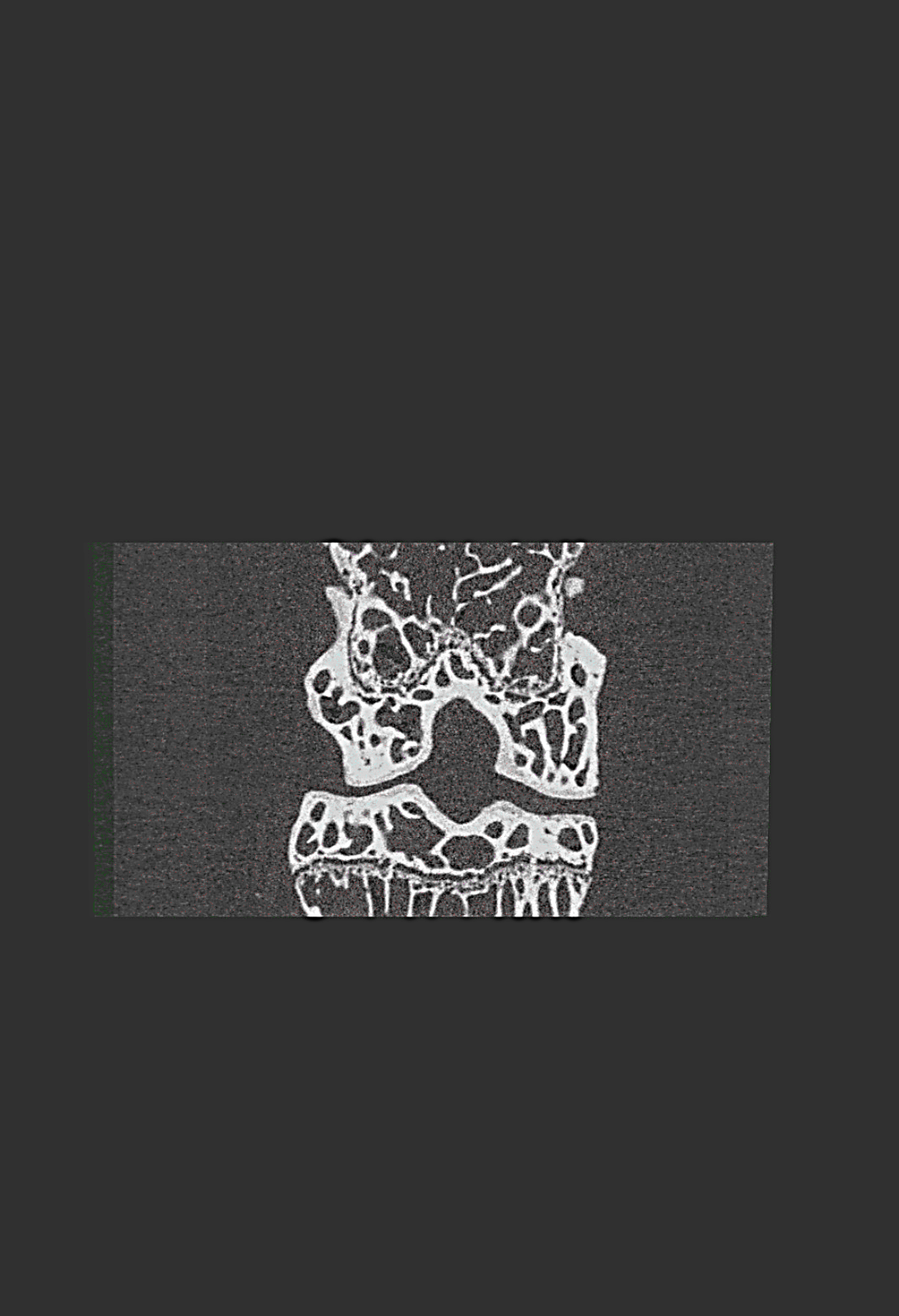

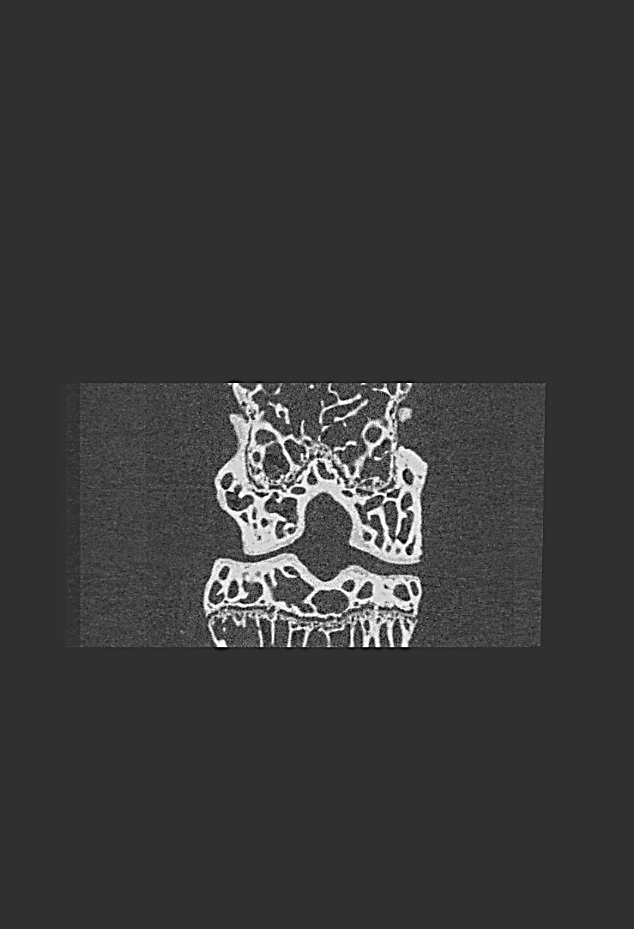

**Sham**

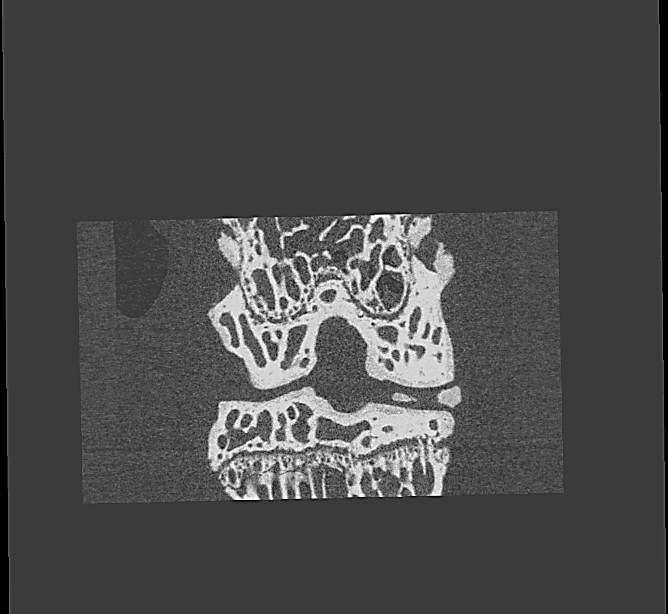

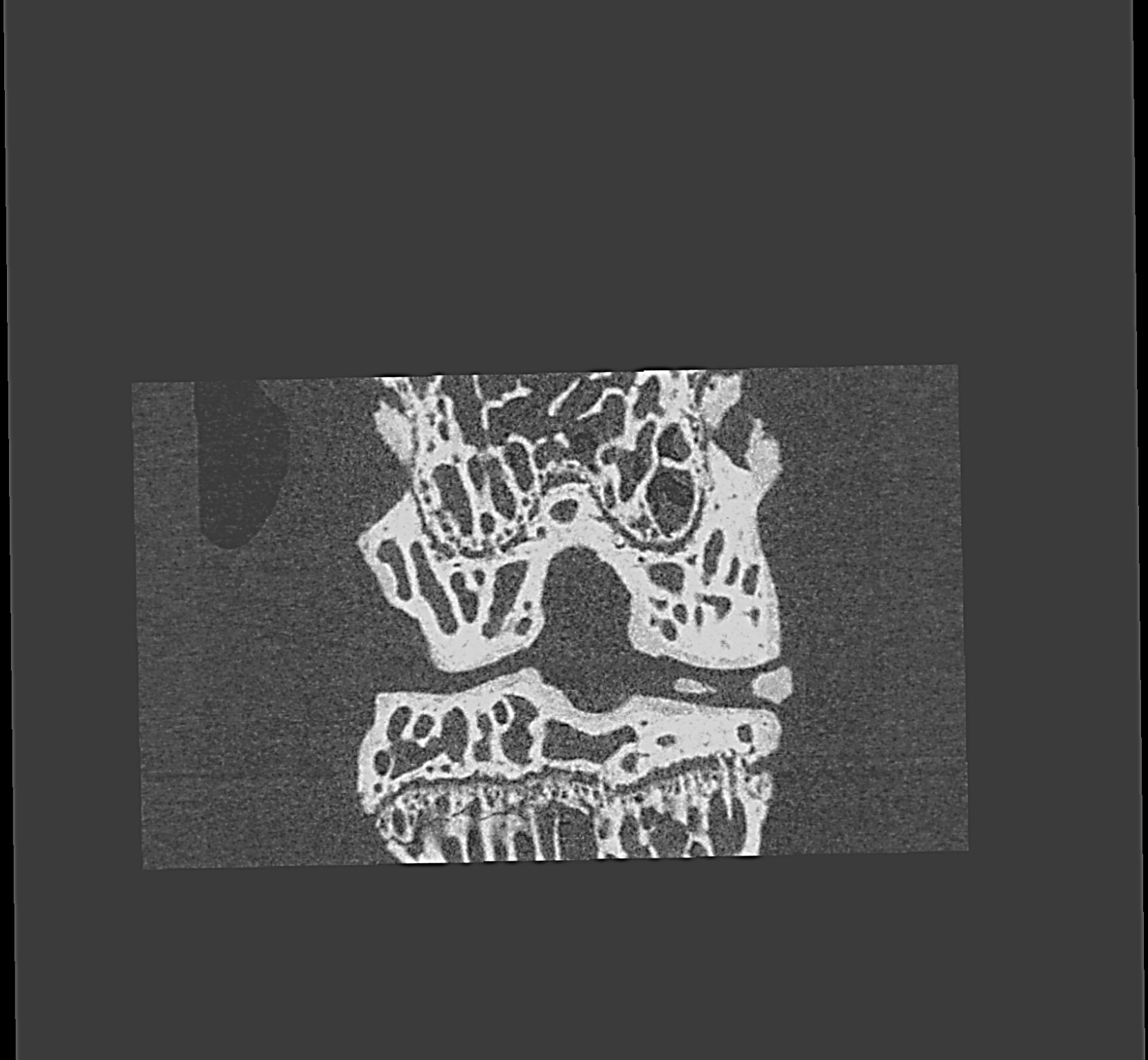

**DMSO**

**TG-18 + L-006235 every 4 week**

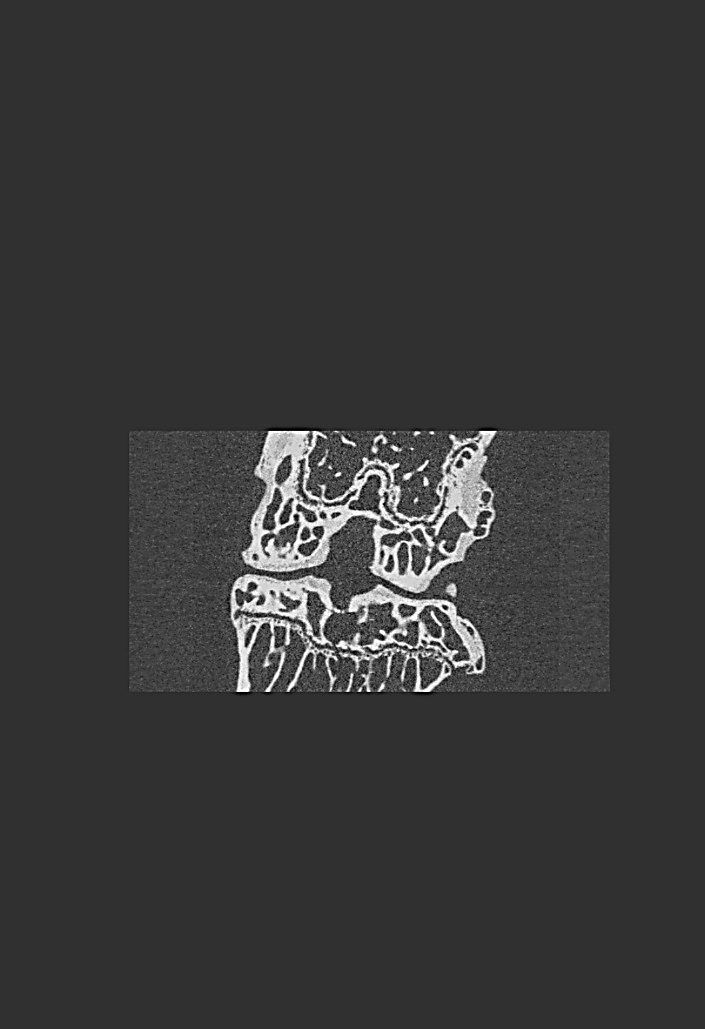

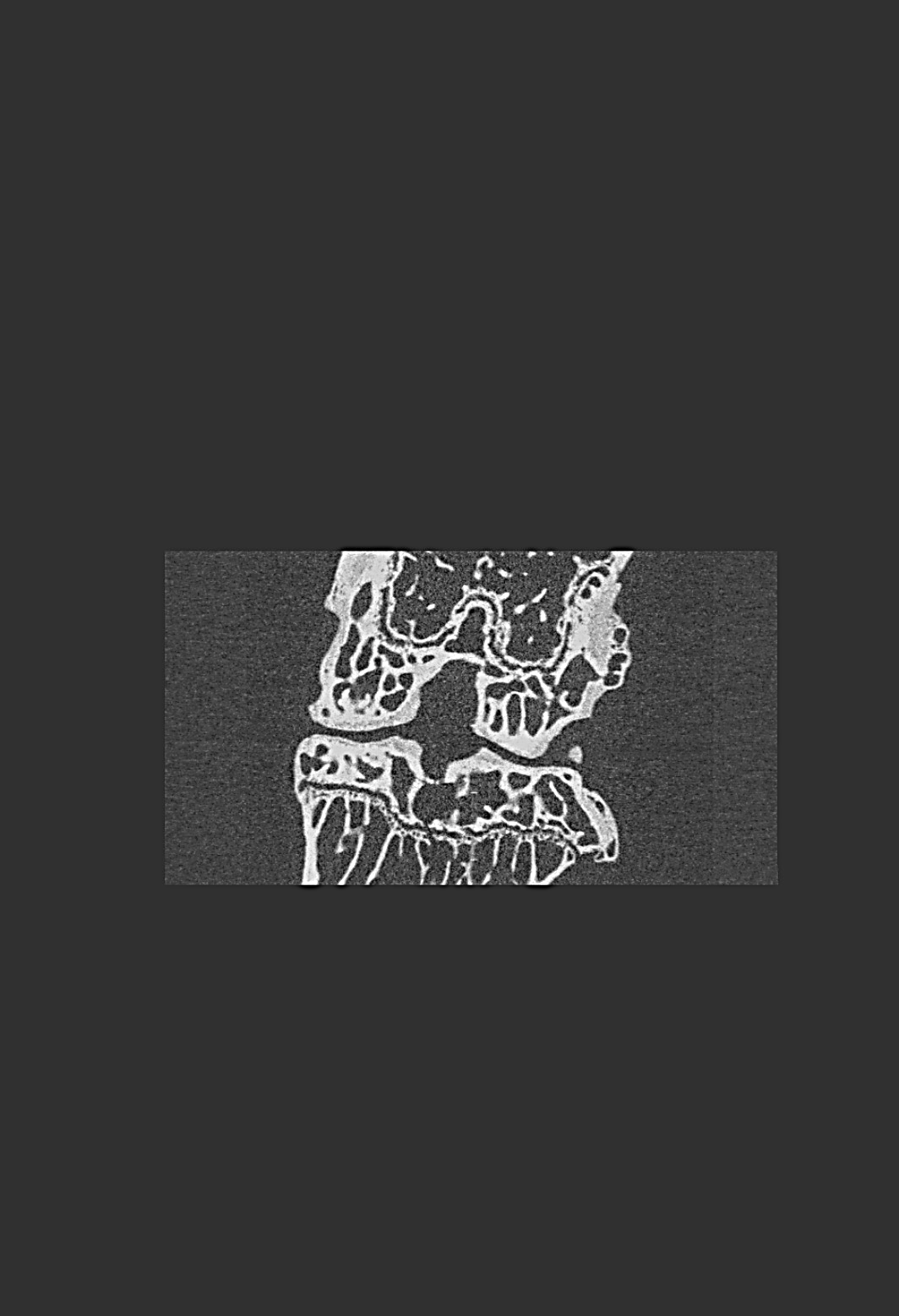

**TG-18 + L-006235 every 2 week**

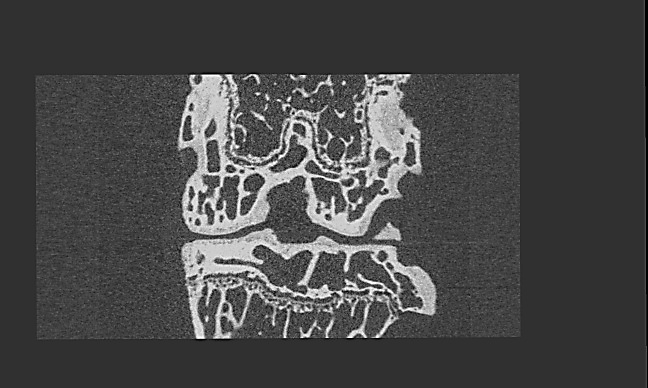

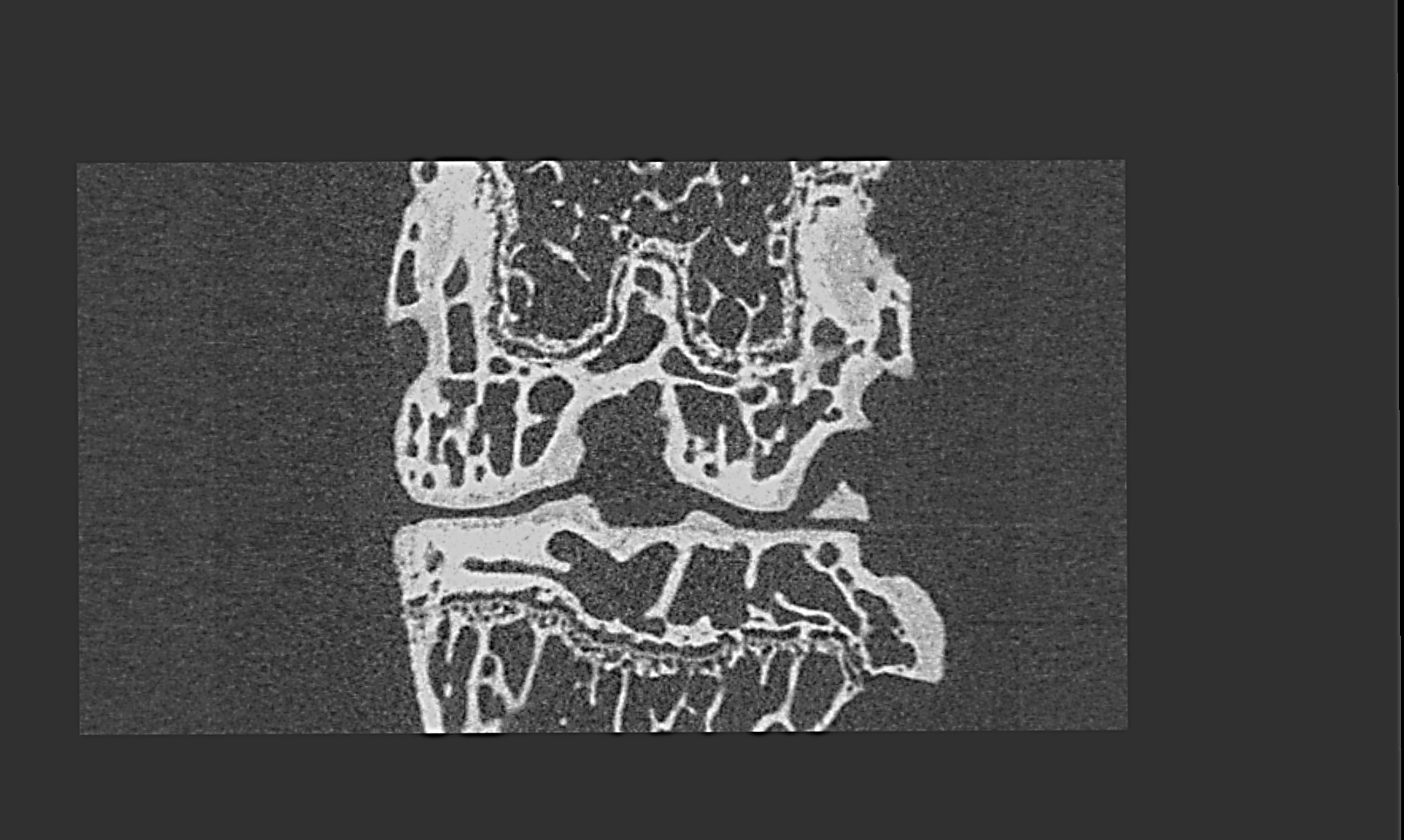

**Sucrose Stearate + L-006235 ( 2 weeks)**

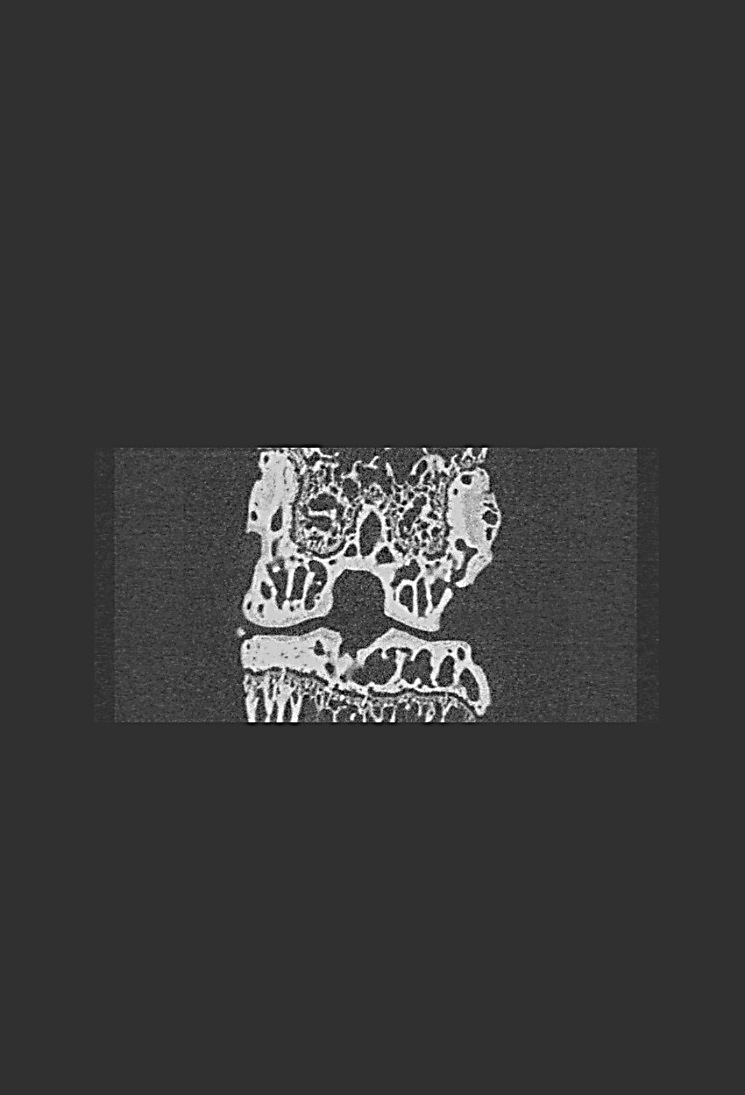

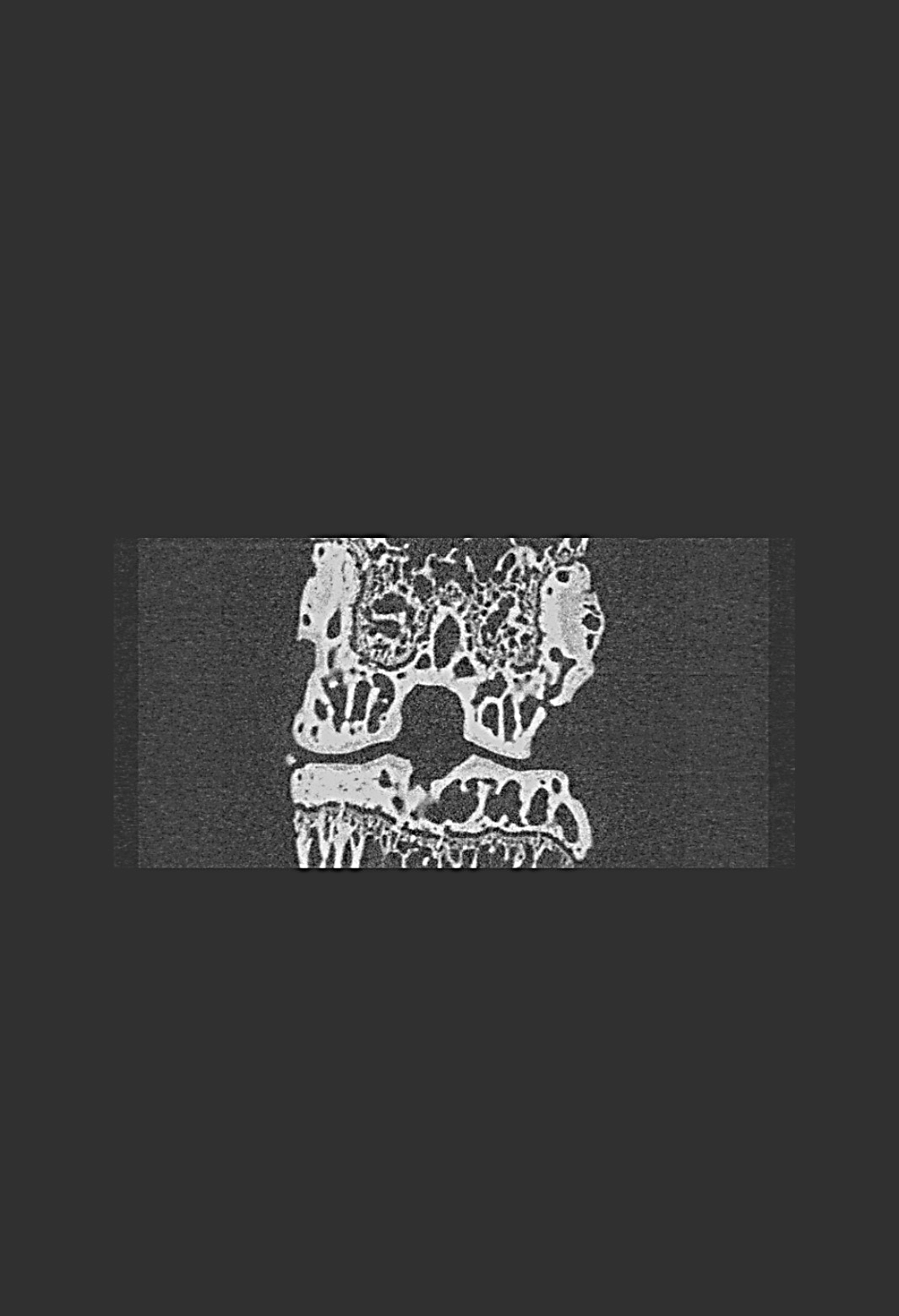

**Sucrose stearate every 4 weeks**

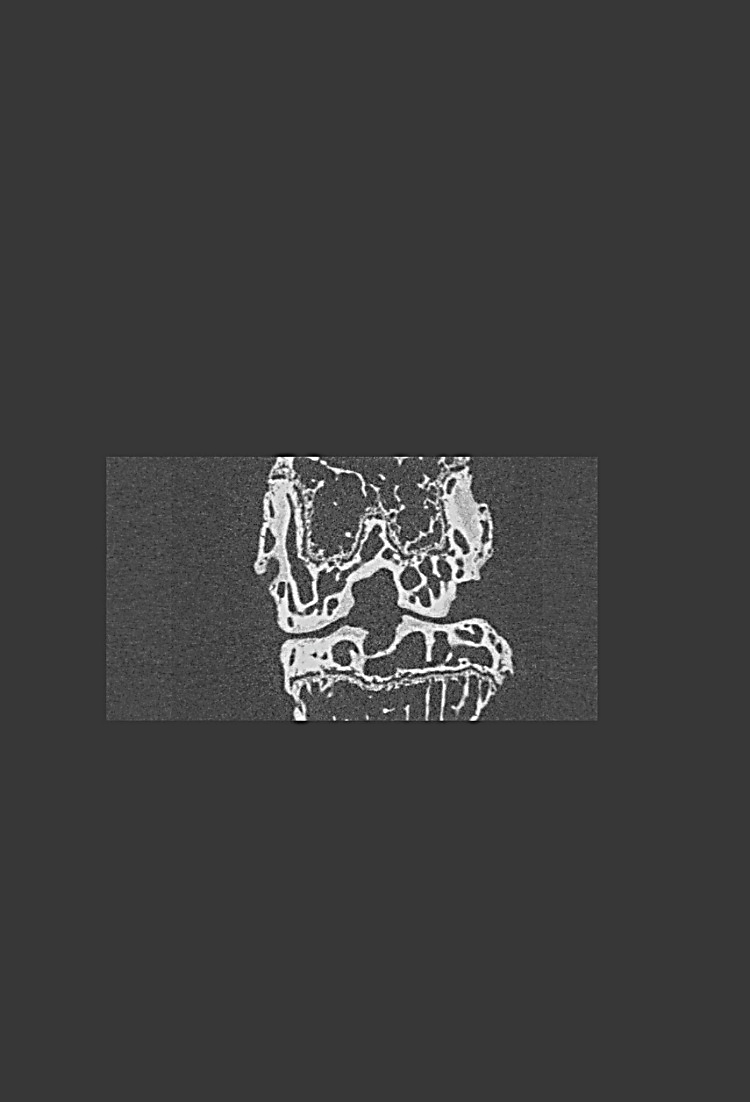

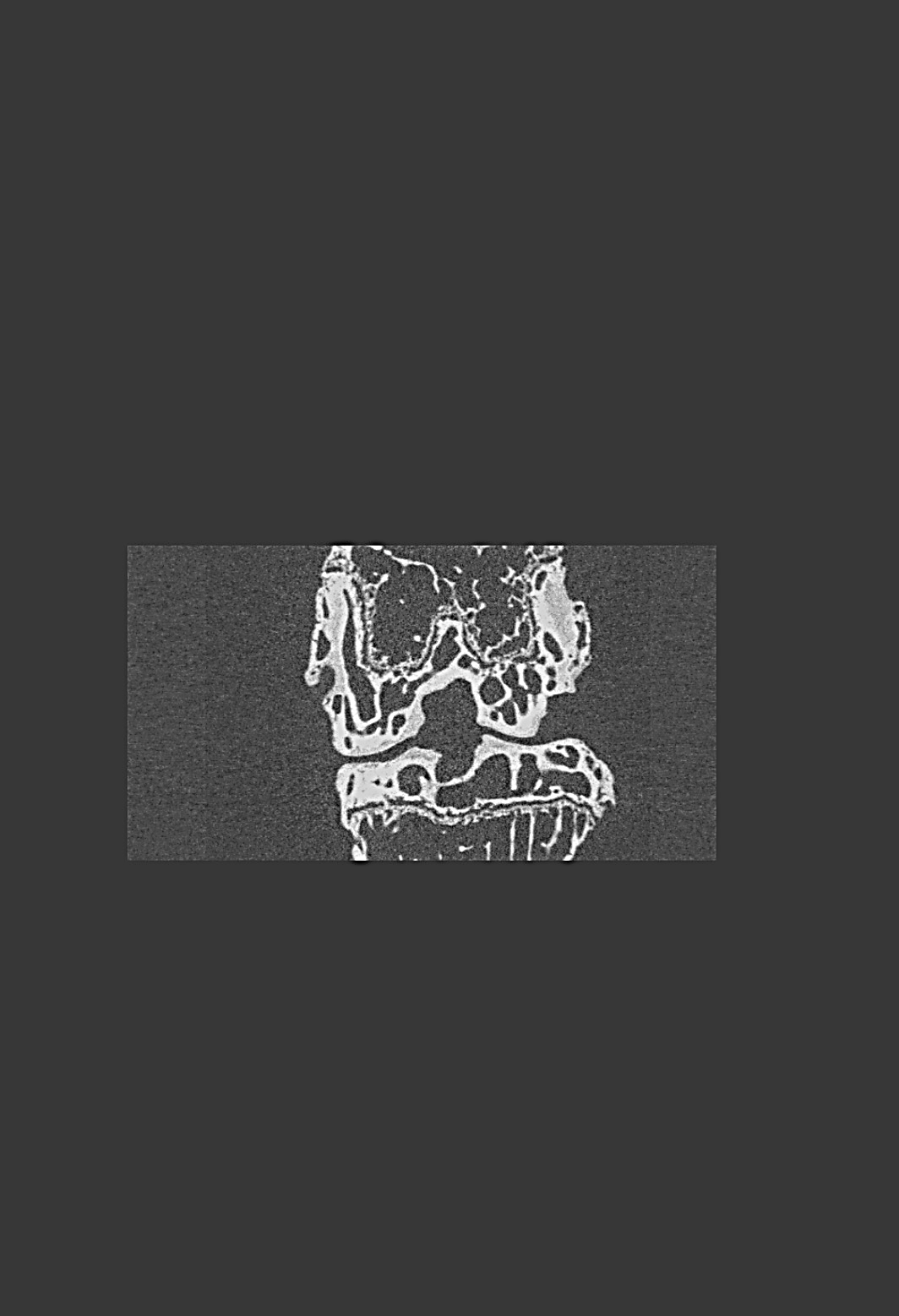

**Healthy**

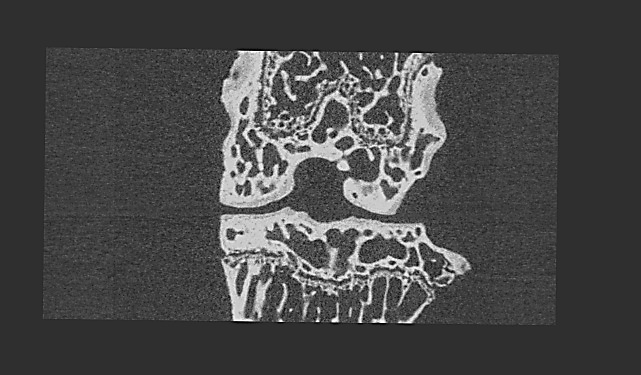

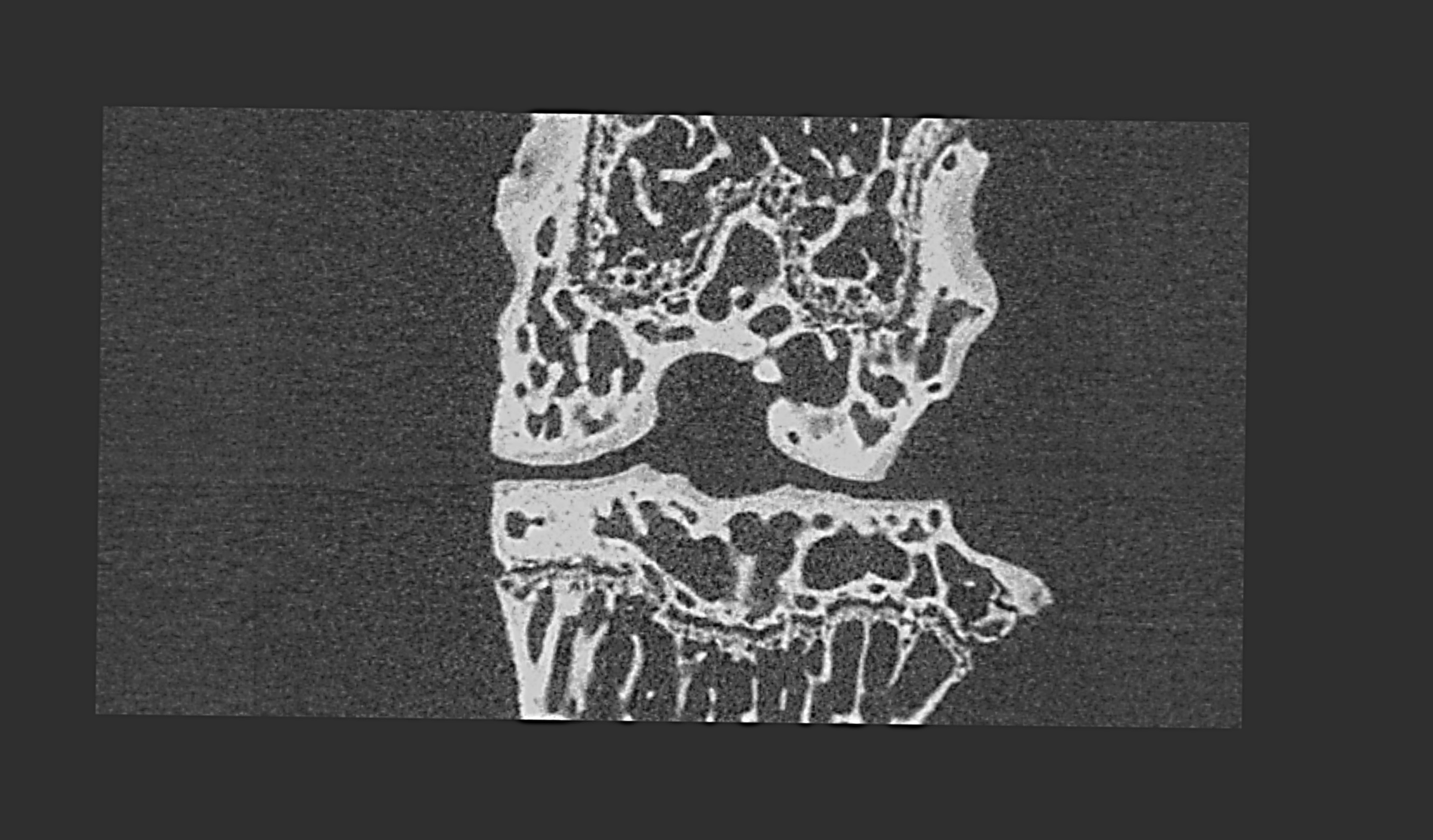
